# Conserved architecture of Tc toxins from human and insect pathogenic bacteria

**DOI:** 10.1101/596536

**Authors:** Franziska Leidreiter, Daniel Roderer, Dominic Meusch, Christos Gatsogiannis, Roland Benz, Stefan Raunser

## Abstract

Tc toxin complexes use a syringe-like mechanism to penetrate the membrane and translocate a toxic enzyme into the host cytosol. They are composed of three components: TcA, TcB and TcC. Until recently, low-resolution structures of TcA from different bacteria suggested that Tc toxins differ considerably in their architecture and possibly in their mechanism of action. Here, we present high-resolution structures and functional studies of five TcAs from different insect and human pathogenic bacteria. Contrary to previous expectations, their overall composition and domain organization is almost identical. The TcAs assemble as a pentamer with a central α-helical channel surrounded by a shell composed of conserved α-helical domains and variable β-sheet domains. Essential structural features, including a conserved trefoil protein knot, are present in all five TcAs, suggesting a common mechanism of action. All TcAs form functional pores and can be combined with TcB-TcC subunits from other species resulting in chimeric holotoxins. We have identified a conserved ionic pair that stabilizes the shell, likely operating as a strong latch that only springs open after the destabilization of other regions. Our results lead to new insights into the architecture and host specificity of the Tc toxin family.

## Introduction

Large multi-subunit Tc toxins are present in a variety of bacterial pathogens. They were first identified in the insect pathogenic bacterium *Photorhabdus luminescens*^1^ and it was shown that they have oral activity against insects^2^. Consequently, the study of Tc toxins has become an important aspect in the search for novel biopesticides as an alternative to *Bacillus thuringiensis* (Bt) toxins^3^. Bt toxins were the first biopesticides that were successfully used in transgenic plants. However, an increasing number of insects have developed resistance against Bt toxins^4^, creating a crucial need for alternatives. Since Tc toxins are also encoded by human pathogenic bacteria like *Yersinia pestis*^5^, *Yersinia pseudotuberculosis*^6^, and *Morganella morganii*^7^ the elucidation of the function and mechanism of action of these toxins is also of high medical relevance.

Tc toxins from *Photorhabdus luminescens* are composed of three subunits: TcA, TcB and TcC^8^. The 1 – 1.4 MDa homopentameric TcA acts as an injecting device, responsible for translocating the actual toxic component into host cells^9,10^. TcA consists of a preformed channel that is surrounded by a shell domain. Putative receptor-binding domains (RBDs) localized at the periphery of the shell possibly interact with receptors on the host cell membrane^11^. pH-induced opening of the shell releases a putative entropic spring that drives the injection of the TcA channel into the membrane^11^. TcB and TcC form a cocoon that harbors the actual toxic component, namely the C-terminal hypervariable region (HVR) of TcC, which is autoproteolytically cleaved^11,12^. Binding of the TcB-TcC cocoon to the TcA channel via a six-bladed β-propeller triggers opening of the cocoon and translocation of the toxic enzyme into the channel^11,13^. During this process, parts of the β-propeller completely unfold and refold into an alternative conformation upon binding^13^. The enzyme passes through a narrow negatively charged constriction site inside the cocoon, most likely acting as an extruder that releases the unfolded protein with its C-terminus first into the translocation channel^13^.

In our recent work, we demonstrated that the transmembrane helices of TcdA1 from *P. luminescens* rearrange once they enter the membrane in order to open the initially closed pore^14^. The linker domain, which is stretched in the prepore state^11^, is folded and tightly packed in a pocket that is formed between the shell and channel domains^14^.

Besides the high-resolution structures of TcdA1 in its prepore and pore state^11,13,14^, only low-resolution structures of two other TcAs have been determined. Whereas the TcA component of YenTCA1A2 from *Yersinia entomophaga*, like TcdA1 from *P. luminescens*, forms a pentameric bell-shaped complex with an inner channel and an outer shell^9^, the structure of XptA1 from *Xenorhabdus nematophila* appears to have a tetrameric cage-like structure with a central cavity^15^. This discrepancy indicates that Tc toxins differ considerably in their architecture and as a consequence, possibly in their mechanism of action.

To better understand the structural variety of Tc toxins and to obtain a holistic view on their architectural organization and function, we determined the structures of TcA components from different insect and human pathogenic bacteria using single particle electron cryo microscopy (cryo-EM). The study included TcdA1 and TcdA4 from *P. luminescens* (Pl-TcdA1 and Pl-TcdA4), XptA1 from *Xenorhabdus nematophila* (Xn-XptA1), TcdA4 from *Morganella morganii* (Mm-TcdA4) and TcaA-TcaB from *Yersinia pseudotuberculosis* (Yp-TcaATcaB). By using our single particle processing software SPHIRE^16^, we obtained resolutions of up to 2.8 Å.

Contrary to previous expectations, the structures revealed that the examined TcAs, including Xn-XptA1, share the same pentameric bottle-shaped structure and that the domain organization of these toxin components is almost identical. Our study demonstrates that functionally crucial, structural features, i.e. the linker domain, the translocation channel and the TcB-binding domain have high structural similarities across all analyzed TcAs.

The main structural differences between the TcAs are found at the periphery of the shell where the RBDs and the neuraminidase-like domain are located. Surprisingly, the bottom of the shell, formed by the neuraminidase-like domain, is not completely closed in Xn-XptA1, Mm-TcdA4 and Yp-TcaATcaB. Interestingly, we identified a novel coiled coil domain in Yp-TcaATcaB, that is not found in any of the other studied TcAs. It reaches out from the shell and interacts with the funnel-shaped TcB-binding domain of TcA. Inversed charges close to the tip of the translocation channel in Xn-XptA1, Mm-TcdA4 and Yp-TcaATcaB indicate a possible difference in protein translocation. Overall, our results suggest that Tc toxins from different organisms share a common architecture and consequently mechanism of action, while the variability in the RBDs enables the targeting of different hosts.

## Results and Discussion

### TcA toxin subunits share a common architecture

In order to exclude that the crystallization buffer or crystal contacts had an influence on our previous crystal structure of Pl-TcdA1, we first determined a 2.8 Å resolution structure of Pl-TcdA1 from *P. luminescens* using cryo-EM and single particle analysis in SPHIRE^16^. The overall structure of Pl-TcdA1 was the same, but the higher resolution, 2.8 Å in comparison to 4 Å, enabled us to improve the atomic model of the complex (Figure 1a, Supplementary Figure 1a, Supplementary Figure 2, Supplementary Video 1).

**Figure 1:**
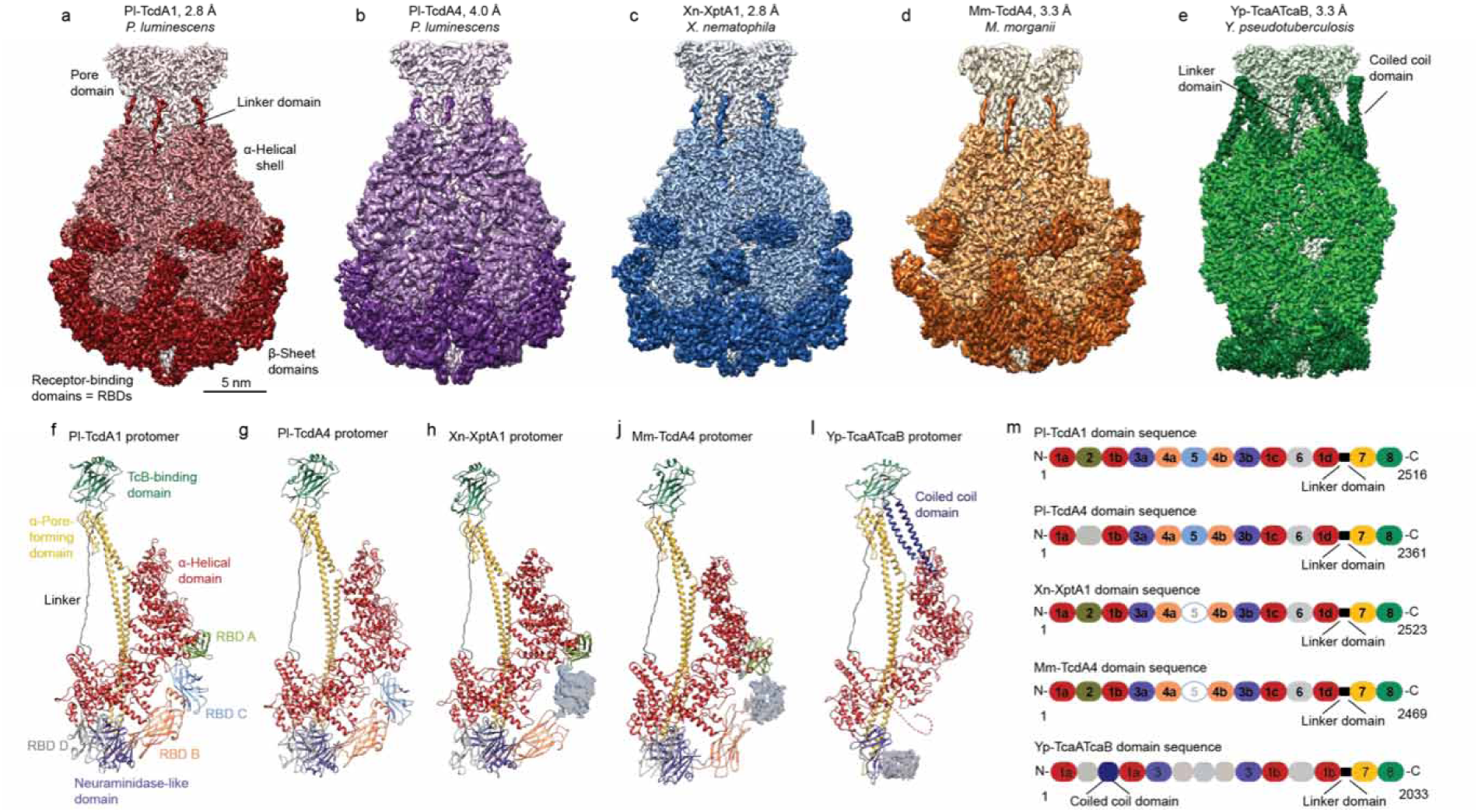
Structures of five TcAs. (**a-e**) Cryo-EM density maps of Pl-TcdA1, Pl-TcdA4, Xn-XptA1, Mm-TcdA4 and Yp-TcaATcaB, respectively, with the average resolutions according to 0.143 FSC. The color gradient from light to dark represents the pore domain with the TcB-binding domain, the α-helical shell, the β-sheet domains and the linker. (**f-k**) Structures of the TcA protomers. Pl-TcdA4 does not contain RBD A. RBD C was not well resolved in Xn-XptA1 and Mm-TcdA4 and is therefore not included in the models. Yp-TcaATcaB does not contain any RBD. The unique coiled coil domain of Yp-TcaATcaB is highlighted in dark blue. 99 residues (amino acids 1140-1239) of the neuraminidase-like domain and the first 57 residues at the N-terminus of Yp-TcaATcaB could not be built. The densities of the domains that could not be built are shown to indicate their location (**h-k**). The N-terminus of Yp-TcaATcaB (residues 1-57) is depicted as red dotted line. (**l**) Domain organization of Pl-TcdA1, Pl-TcdA4, Xn-XptA1, Mm-TcdA4 and Yp-TcaATcaB. 1 = helical shell, 2 = RBD A, 3 = neuraminidase-like domain, 4 = RBD B, 5 = RBD C, 6 = RBD D, 7 = channel, 8 = TcB-binding domain and the linker domain in black. Domains that could not be built are shown as dashed circles and domains that are missing in the sequence are faded.

To compare the structure of Pl-TcdA1 with the structure of other TcA complexes, we heterologously expressed Pl-TcdA4, Xn-XptA1, Mm-TcdA4 and Yp-TcaATcaB in *E. coli* and purified the proteins. While the TcAs from *P. luminescens*, *X. nematophila* and *M. morganii* are single proteins, Yp-TcaATcaB is composed of two subunits (TcaA and TcaB). TcaA forms the α-helical shell and TcaB comprises the pore-forming domain as well as the bottom part of the toxin (Supplementary Figure 1f). In order to obtain complete complexes, Yp-TcaATcaB was expressed as a fusion construct (Methods).

We then determined the near-atomic structures of the different TcA complexes using cryo-EM and single particle analysis yielding resolutions of 4.0 Å (Pl-TcdA4), 2.8 Å (Xn-XptA1), 3.3 Å (Mm-TcdA4 and Yp-TcaATcaB) (Figure 1b-e, Supplementary Figure 1b-e, Supplementary Figure 2, Supplementary Figure 3, Supplementary Video 1). The cryo-EM structures allowed us to build atomic models of the complexes revealing that all five studied TcAs share the same overall architecture. They all have a pentameric bell-shaped structure consisting of a central pore-forming domain surrounded by an outer shell, which is made up of an α-helical part as well as β-sheet domains. The linker domain, which connects the pore-forming domain with the outer shell, is present in all examined TcAs (Figure 1a-e).

Importantly, in contrast to the previously reported low-resolution structure of Xn-XptA1 which suggested a tetramer with a cage-like structure^15^, our structure of Xn-XptA1 clearly reveals that it is a pentameric complex with a bell-shaped appearance similar to Pl-TcdA1 from *P. luminescens*. The relatively high amino acid sequence identity of roughly 46.7% between Xn-XptA1 and Pl-TcdA1 supports our results.

### Differences in the shell domain

In all TcAs, the inner scaffold of the shell is composed of an α-helical domain (Figure 1l). It can be divided into a large and a small lobe that are arranged perpendicular to each other forming an L-shape (Figure 2a). The small lobe can be further subdivided into two pseudo-2-fold-symmetrical subunits in an X-shaped structure (Figure 2a). The large lobe contains two different pseudo repeats (Figure 2b,c), which are composed of several helix-loop-helix motifs. Both pseudo repeats have the same fold and organization for all five TcAs. However, the pseudo repeat 2 domain of Yp-TcaATcaB contains two additional insertions, namely an enlarged loop and a coiled coil domain (Figure 1e,k, Figure 2d). The coiled coil domain reaches out to the TcB-binding domain (Supplementary Figure 1f), thus connecting the two subunits TcaA and TcaB. The coiled coil domain probably increases the stability of the complex, which, in contrast to the other TcAs, does not contain any RBDs (see below).

**Figure 2:**
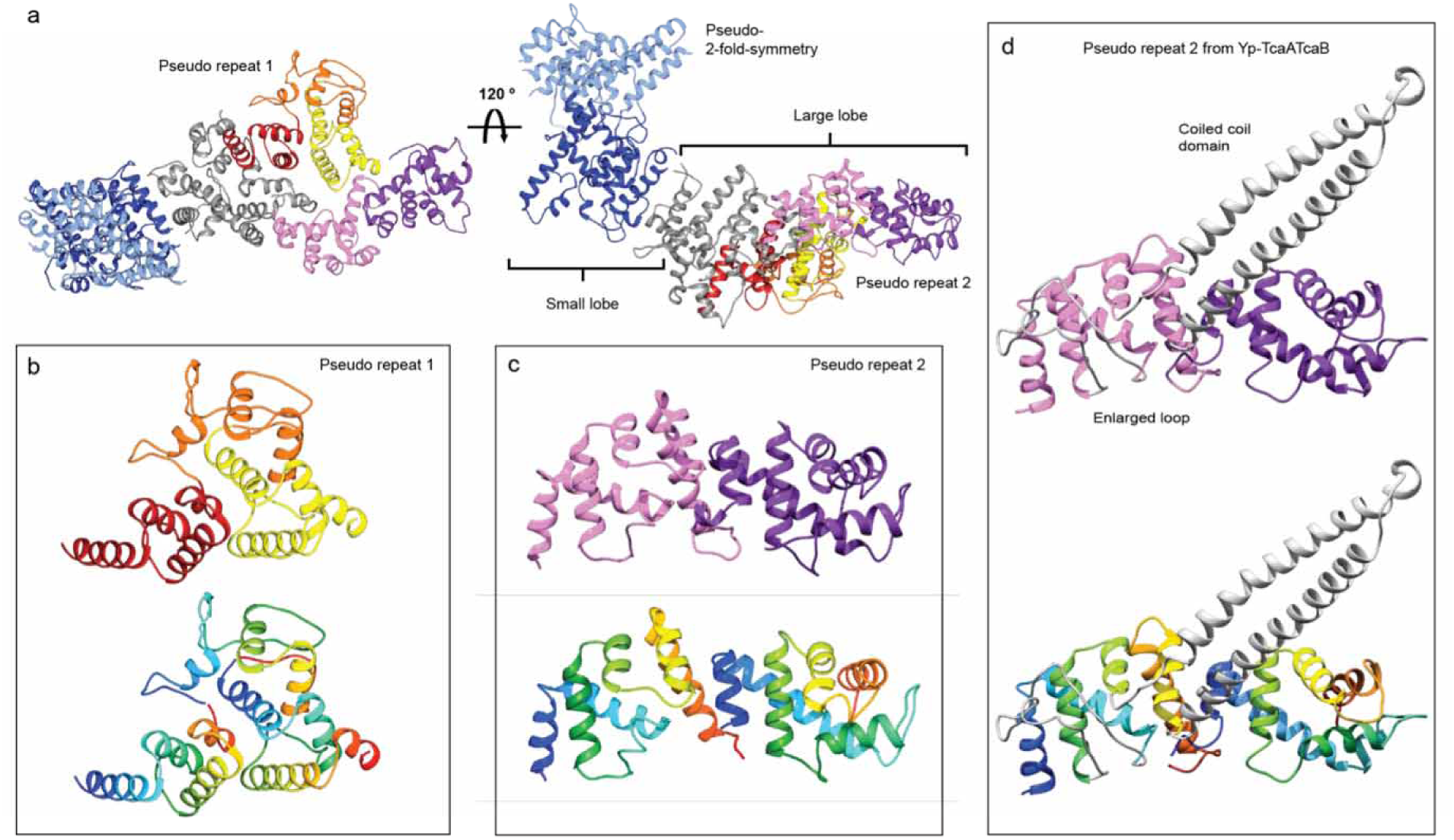
Organization of the α-helical shell. (**a**) The α-helical shell of a TcA protomer shown for Pl-TcdA1 can be divided into a small lobe (amino acids 1-160, 964-1090, 1608-1632, 1762-1972) and a large lobe (amino acids 161-297, 434-963). The small lobe has a pseudo-2-fold symmetry (light and dark blue), resulting in an X-shaped structure. The large lobe contains two pseudo repeats; pseudo repeat 1 is shown in red, orange, and yellow and pseudo repeat 2 is shown in rose and magenta. (**b**) The three repeating subdomains of pseudo repeat 1 are depicted in red, orange and yellow in the upper panel as well as in rainbow colors (colored from blue to red from N- to C-terminus) in the lower panel. (**c**) The two repeating subdomains of pseudo repeat 2 are depicted in rose and magenta in the upper panel as well as in rainbow colors in the lower panel. (**d**) The pseudo repeat 2 in Yp-TcaATcaB shows the same overall fold, except for the insertion of the coiled coil domain as well as an enlarged loop (both in grey).

The β-sheet domains of the shell are less conserved than its α-helical domain (Supplementary Figure 1g). The main differences are mostly located at the RBDs (Figure 1a-e). Pl-TcdA1 has four RBDs and a neuraminidase-like domain; RBD A is inserted in the large lobe, and RBD B, C and D as well as the neuraminidase-like domain are inserted in the small lobe (Figure 3d). Whereas Xn-XptA1 and Mm-TcdA4 have the same number of RBDs as Pl-TcdA1, Pl-TcdA4 misses RBD A, resulting in a slimmer shaped molecule (Figure 1). Yp-TcaATcaB does not contain any RBD, only the neuraminidase-like domain (Figure 1e,l). This leads to a more distinct and slimmer shape of the shell domain of Yp-TcaATcaB. The N-terminal domain of TcaA (residues 1-57) and a larger peripheral domain of the neuraminidase-like domain (residues 1140-1239) were not resolved in Yp-TcaATcaB. Due to the missing RBDs, these regions are probably more flexible than in the other TcAs.

**Figure 3:**
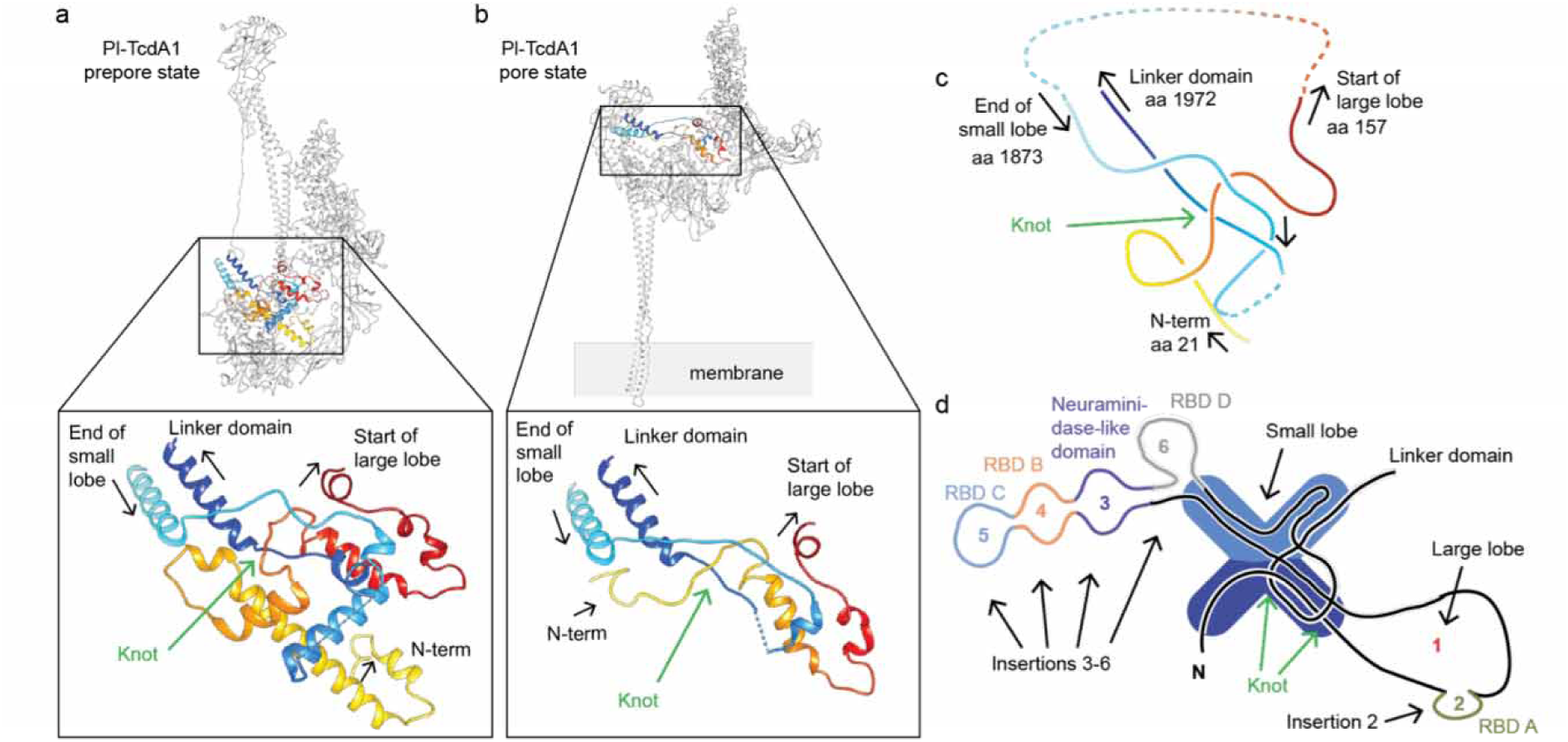
A trefoil 3_1_ protein knot is present in all TcAs. (**a**) A trefoil 3_1_ protein knot is present in all five TcA prepores. The knot structure in a Pl-TcdA1 prepore protomer and a close-up view is presented. The polypeptide chain is highlighted in rainbow colors. The dashed line indicates missing residues (amino acids 1933 – 1938). (**b**) The protein knot is also present in the Pl-TcdA1 protomer in the pore state (pdb ID 5LKH and 5LKI). The dashed line indicates missing residues (amino acids 1914 – 1946). (**c**) Simplified structure of the protein knot of Pl-TcdA1. The arrows indicate the direction of sequence and the amino acid numbering corresponds to Pl-TcdA1. (**d**) Scheme of the shell organization of a TcA protomer showing the location of the knot in the small lobe. RBD A-D and the neuraminidase-like domain are inserted in the main sequence of the big and small lobe.

### A conserved 3_1_ trefoil protein knot stabilizes the basis of the linker

The shell of all five TcAs contains a 3_1_ trefoil^17^ protein knot (Figure 3a, Supplementary Figure 4a-d, Supplementary Figure 5 and Supplementary Video 2). The knot tightly connects the N-terminal part of the protein (residues 21-157 in Pl-TcdA1) with the last domain of the α-helical shell before the linker (residues 1873-1972 in Pl-TcdA1) (Figure 3c,d and Supplementary Figure 5b). The protein knot is also present in the pore state of Pl-TcdA1^14^, indicating that it does untangle during prepore-to-pore transition (Figure 3b).

A 3_1_ trefoil protein knot is the most simple and common knot type^18^. It mostly occurs in smaller proteins or enzymes such as methyltransferases and carbonic anhydrases^19^. With ∼280 kDa (mass of Pl-TcdA1 monomer), TcA is, to our knowledge, by far the largest protein for which such a knot has been observed. The folding mechanism of a knotted protein has been shown to be quite complex and the folding rate of knotted proteins is decreased^18,20^. These findings suggest that it is rather surprising that the complex TcA structure contains a knot.

In many cases, the function of protein knots is unknown^21^. Several studies, however, indicate that a protein knot has a stabilizing effect on proteins^22^, particularly during mechanical stress and conformational changes^23^. In the case of Tc toxins, the knot is indeed strengthening the basis of the linker, which is especially put under strain during the prepore-to-pore transition (Figure 3a,b, Supplementary Figure 5b and Supplementary Video 2). Consequently, the protein knot might also stabilize the stretched linker domain in the prepore state of TcA. Taken together, we conclude that the protein knot in TcA is an important conserved structural feature and propose that it is essential for the structural stability and function of all Tc toxins.

### Differences in the pore domain

Between all TcAs, the pore domain is the part of the protein with the highest conservation (Supplementary Figure 1g, Supplementary Figure 6d) and the highest structural similarity (Supplementary Figure 1h). In addition, it is the among the best resolved regions in all TcA reconstructions (Supplementary Figure 2e,k,p, Supplementary Figure 3e,k). The differences between Pl-TcdA4 and Pl-TcdA1 are neglectable in the pore domain. As such, we will only compare Xn-XptA1, Mm-TcdA4, Yp-TcaATcaB and Pl-TcdA1. The hydrophobic surface properties of the channel exterior are comparable in all TcAs (Supplementary Figure 6a). The tip of their channels (5 nm) is highly hydrophobic, indicating that they have a similarly sized transmembrane domain (Supplementary Figure 6a). Interestingly, whereas the hydrophobic loops at the tip of the transmembrane domain are highly conserved, the residue residing in the center of each loop is variable, suggesting different interactions with the host membrane (Supplementary Figure 6d,e). The surface potentials are similar in Pl-TcdA1, Xn-XptA1 and Mm-TcdA4 with a patch of positive charges at the upper part and a patch of negative charges at the lower part of the α-pore forming domain. The channel of Yp-TcaATcaB, however, is more negatively charged at its outside and misses the prominent stretch of positive charges (Supplementary Figure 6b).

The lumen of the channel of Pl-TcdA1 contains several discrete bands of negative electrostatic potential^11,14^ (Figure 4a). Therefore, Pl-TcdA1 is selective to cations if reconstituted in membranes without TcB-TcC^11,24^. The pattern of negative charges is also found in the lumen of the other TcA channels. However, the lower one to two bands at their tip are positively charged instead (Figure 4a).

**Figure 4:**
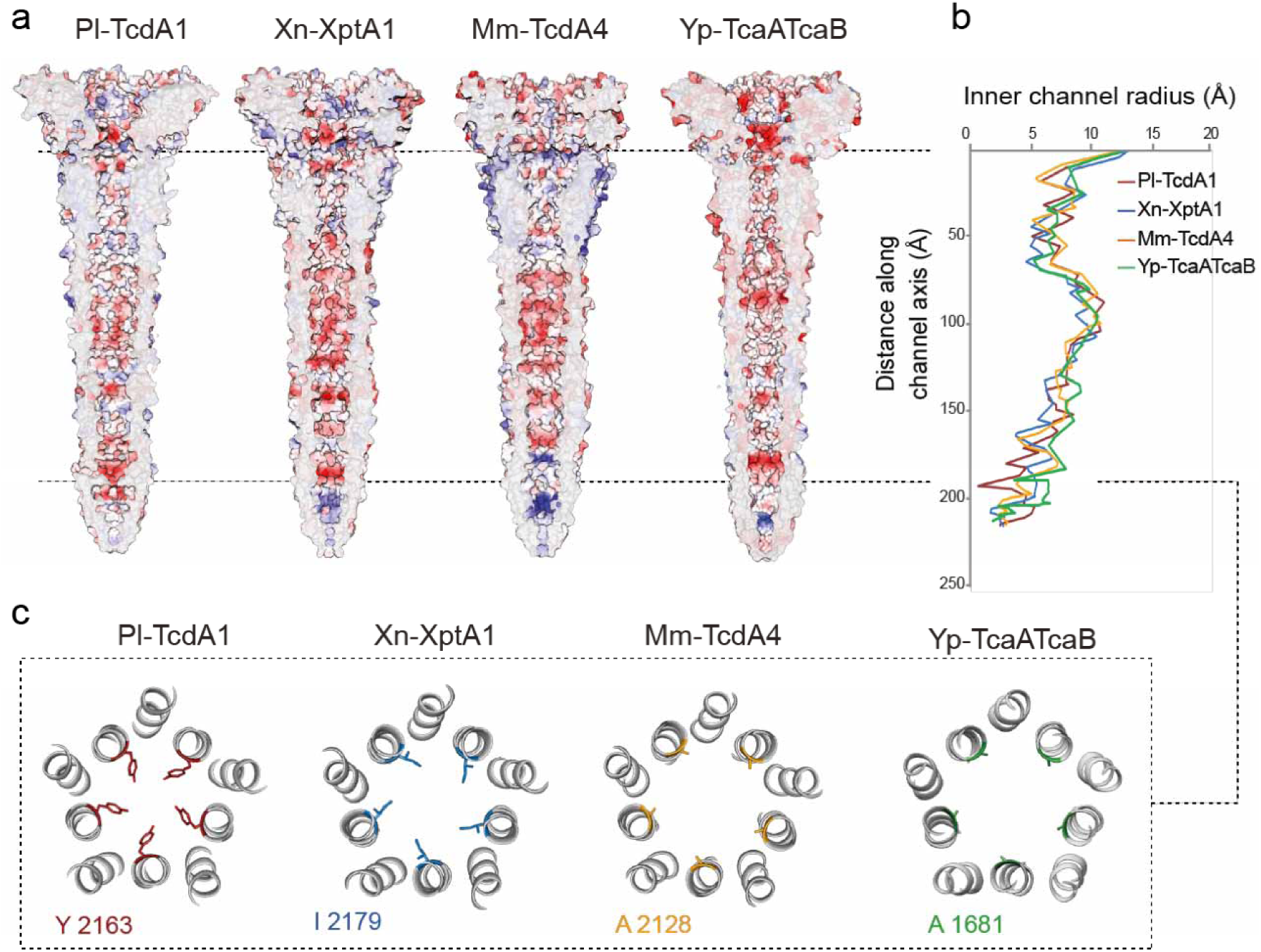
Comparison of the TcA channels. (**a**) Cross-sections of the TcA channels, demonstrating the electrostatic Coulomb potential in the channel lumen of Pl-TcdA1, Xn-XptA1, Mm-TcdA4 and Yp-TcaATcaB at pH 7. Positively charged (14 kcal/mol) and negatively charged (−14 kcal/mol) residues are colored in blue and red, respectively. (**b**) Graph depicting the inner channel radius of Pl-TcdA1 (red), Xn-XptA1 (blue), Mm-TcdA4 (orange) and Yp-TcaATcaB (green). At the narrowest position of the Pl-TcdA1 prepore (Y 2163, indicated by the dashed line), the diameter of the channel is 3.9 Å. In Xn-XptA1, Mm-TcdA4 and Yp-TcaATcaB, the channel diameter reaches only 8.2, 7.8 and 8.4 Å at the narrowest part, respectively. (**c**) Slice through the channel at the position of the Pl-TcdA1 channel constriction (Y2163).

The channel lumen shows a comparable diameter profile for the four TcAs (Figure 4b). Starting with a large diameter of around 26 Å for all TcAs at the TcB-interacting domain, the channel radius is decreased along the transport direction of the toxin. Between 40 – 65 Å from the channel entry, all TcA channels are narrowed to a diameter of around 10 Å. Afterwards, the channel widens up and reaches a diameter of up to 20 Å. At this position, there is an accumulation of hydrophobicity in all TcA channels (Supplementary Figure 6c).

A major difference between the different TcAs can be found at the tip of the channel. Here, Y2163 in Pl-TcdA1 forms a constriction site with a diameter of 3.9 Å^11^ (Figure 4c) that opens after membrane insertion^13^. Xn-XptA1, Mm-TcdA4 and Yp-TcaATcaB have smaller residues (I2179, A2128 and A1681, respectively) at this position, resulting in wider diameters with minima of 8.6, 8.9 and 8.4 Å, respectively (Figure 4c). The tip is closed in all TcAs in the prepore conformation (Figure 4a).

To understand whether the minor differences in the channel properties have an influence on its function, we reconstituted TcA into black lipid bilayers and measured the conductance of the channel. All TcAs formed ion-permeable pores within a pH range of pH 4 – 11 (Supplementary Figure 7). In accordance with our previous work, Pl-TcdA1 showed a higher pore-forming activity at extreme pH values compared to pH 6^11^ which goes along with the results obtained for Xn-XptA1 and Mm-TcdA4. Interestingly, at pH 6, Yp-TcaATcaB incorporated more readily into the membrane than the other TcAs. In addition, Yp-TcaATcaB was in general less stable at more extreme pH values (Supplementary Figure 8), which is also reflected in noisier single channel currents at pH 4 and 11 (Supplementary Figure 7d).

While the conductance of Mm-TcdA4 is similar to Pl-TcdA1, the mean single channel conductance values of Xn-XptA1 and Yp-TcaATcaB are 100 – 150 pS lower, independent of the pH. Since the diameter of all TcA channels is comparable (Figure 4b), their charge distributions are likely responsible for this difference (Figure 4a).

### The conservation of the TcB-binding domain allows the formation of chimeric holotoxins

The TcB-binding domain is crucial for the assembly of the holotoxin complex and is conserved among the four TcAs. Accordingly, the TcB-binding domains of TcAs from four different organisms (Pl-TcdA1, Xn-XptA1, Mm-TcdA4 and Yp-TcaATcaB) practically share identical folds (Supplementary Figure 9a). The lowest variance is found in the central β-sheet region, both with respect to amino acid conservation and RMSD (Supplementary Figure 9b,c). Outward-facing loops are less conserved and differ slightly in their size. In the central β-sheets, however, almost all residues are identical in the four TcAs (Supplementary Figure 9d).

The identical fold and high sequence conservation together with the high conservation of the β-propeller of TcB^13^ suggest that it is possible to form chimeric holotoxins by combining TcB-TcC complexes with TcA from different bacteria. Indeed, when TcdB2 and TccC3 from *P. luminescens* is co-expressed with XptA2 from *X. nematophila*, a hybrid toxin complex is formed that is highly active against insects^25^.

To examine whether TcB-TcC from *P. luminescens* (Pl-TcdB2-TccC3) can also be combined with other TcAs, we incubated it with all four TcAs. Using negative stain EM, we observed holotoxin complexes for all combinations tested (Figure 5a-d). In the case of Mm-TcdA4 and Yp-TcaATcaB, however, some TcAs were still present that did not interact with TcB-TcC (Figure 5c,d). Expectedly, the measured affinities between Pl-TcdB2-TccC3 and Pl-TcdA1 (wild type complex), Yp-TcaATcaB and Xn-XptA1 were very high (K_D_ = 0.63 ± 0.03 nM, K_D_ = 0.95 ± 0.02 nM, and K_D_ = 1.99 ± 0.15 nM, respectively) (Supplementary Figure 10). In line with the EM results, the affinity between Pl-TcdB2-TccC3 and Mm-TcdA4 was an order of magnitude lower (K_D_ = 11.3 ± 0.2 nM) (Figure 5c,e). All chimeric holotoxins were fully functional and active against HEK293T cells (Supplementary Figure 11). However, the hybrid toxins were less effective than the native holotoxin complex formed by Pl-TcdA1 and Pl-TcdB2-TccC3 (Supplementary Figure 11c-e).

**Figure 5:**
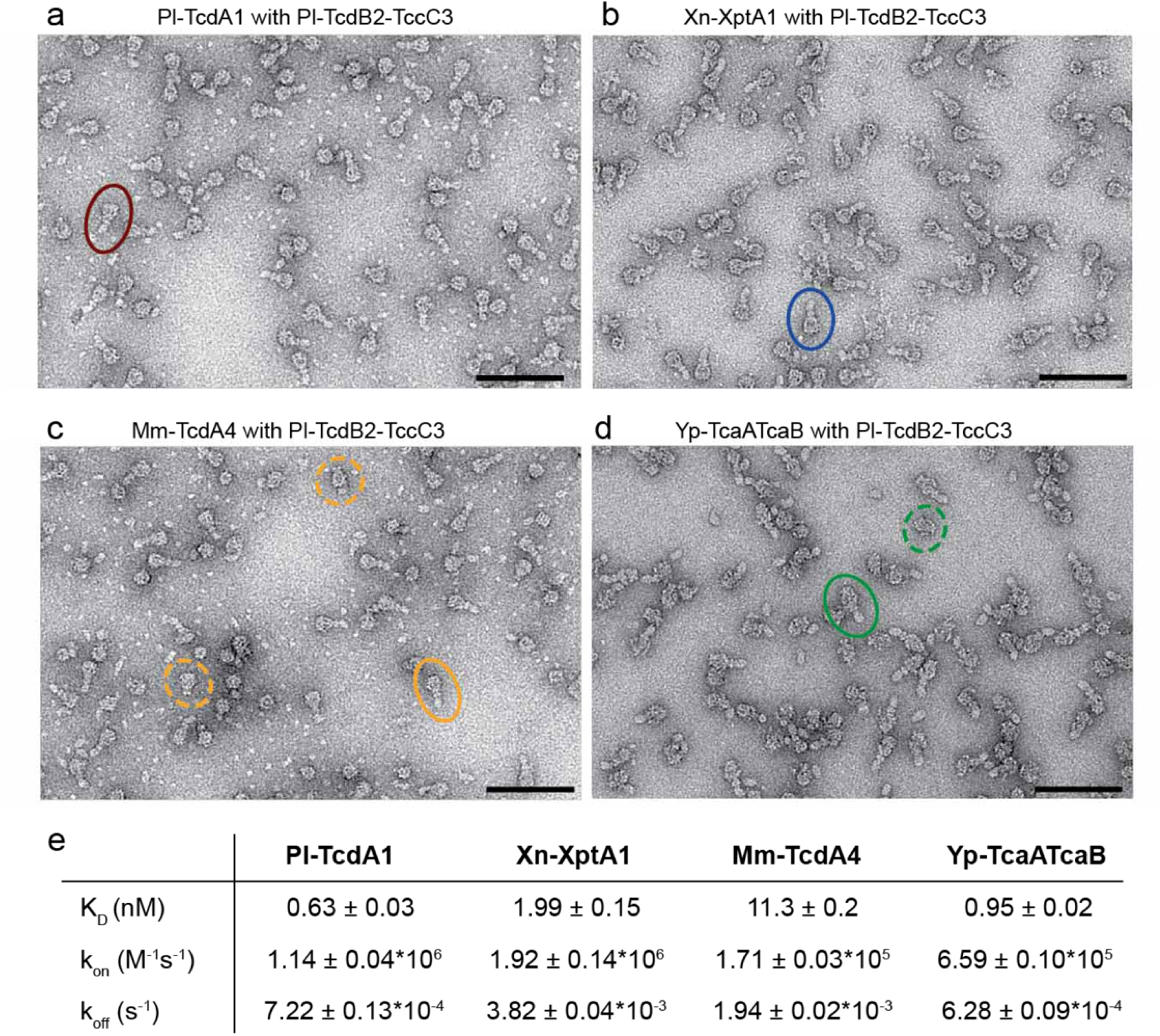
Formation of chimeric holotoxins. (**a-d**) Negative stain electron micrographs after complex formation of different TcAs with TcdB2-TccC3 from *P. luminescens* and size exclusion chromatography. For each complex a holotoxin particle is highlighted by circles. For Mm-TcdA4 and Yp-TcaATcaB, unbound TcAs are marked with dashed circles. Scale bar, 100 nm. (**e**) Table with the measured affinities for the chimeric complexes by biolayer interferometry including the dissociation constant (K_D_) as well as the on- and off-rate of complex formation (k_on_ and k_off_, respectively). A global fit according to a 1:1 binding model was applied, including 6-7 individual curves. The obtained parameters are the mean value ± the error of the fit. See also Supplementary Figure 10.

Altogether, our results demonstrate that functional chimeric holotoxins can be assembled from different Tc subunits. This allows to combine different host specificities with different enzymatic activities and therefore broadens the spectrum of the potential application of Tc toxins as biopesticides.

### The electrostatic lock of the shell domain

The neuraminidase-like domain closes the shell at the bottom of Pl-TcdA1^11^ (Supplementary Figure 1a, Figure 6a-c). Based on the charge distribution in this region, we previously proposed that this domain functions as an electrostatic lock^11^, which is closed at neutral pH and opens at high or low pH due to the repulsion of charged amino acids, triggering the prepore-to-pore transition of Pl-TcdA1^11^.

**Figure 6:**
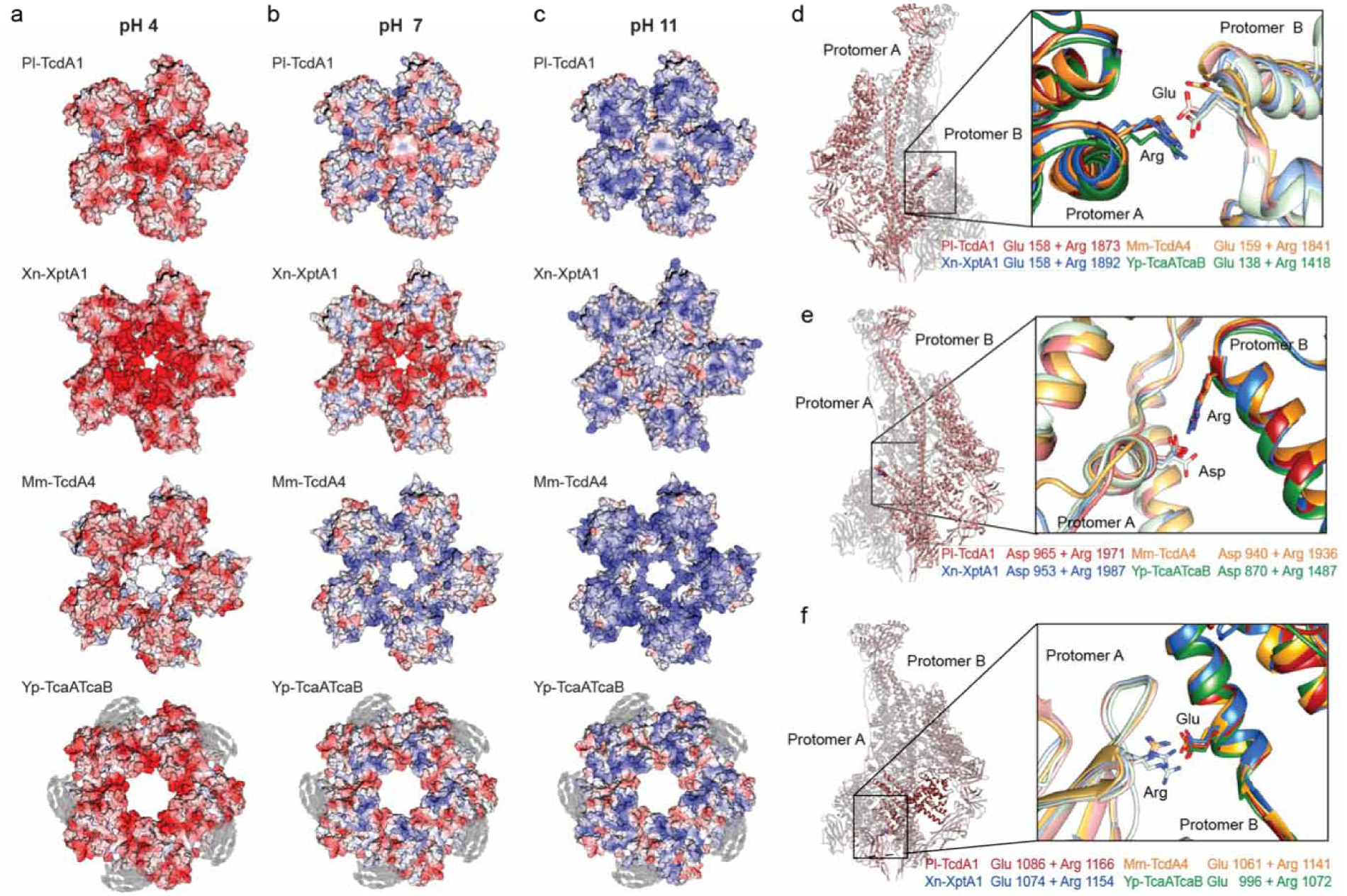
Electrostatic potential of the neuraminidase-like domain and conserved ionic interactions in the shell of TcAs. (**a-c**) Surface electrostatic Coulomb potential of the neuraminidase-like domain at different pH values, viewed from the bottom of TcA. Positively charged (14 kcal/mol) and negatively charged (−14 kcal/mol) residues are colored in blue and red, respectively. Surface electrostatic Coulomb potential at pH 4 (**a**), pH 7 (**b**), and pH 11 (**c**) are shown. The cryo-EM density map of the 99 residues, that could not be built in Yp-TcaATcaB, is depicted in grey. (**d-f**) Conserved ionic interactions between two protomers in Pl-TcdA1, Xn-XptA1, Mm-TcdA4 and Yp-TcaATcaB. The left panel shows two Pl-TcdA1 protomers indicating the different interaction sites and the right panel presents close-up views of each interaction for Pl-TcdA1 (red), Xn-XptA1 (blue), Mm-TcdA4 (orange) and Yp-TcaATcaB (green). (**d**) Interaction of a glutamate (protomer A) with an arginine (protomer B). The residue distance is 3.8 Å in the prepore and 9.2 Å in the pore state of Pl-TcdA1. (**e**) Interaction of an aspartate (protomer A) with an arginine (protomer B). The residue distance is 3.5 Å in the prepore and 5.8 Å in the pore state of Pl-TcdA1. The interacting residues (**d**) and (**e**) belong to the α-helical shell domain. (**f**) Interaction of an arginine (neuraminidase-like domain of protomer A) with a glutamate (small lobe of protomer B). The residue distance is 3.9 Å in the prepore and 25 Å in the pore state of Pl-TcdA1.

To find out whether this is a general feature of TcAs, we compared their neuraminidase-like domains. The fold of the neuraminidase-like domain from different TcAs is almost identical (Supplementary Figure 12b). Only the position of the second subdomain (residues 1140 – 1239) differs in Yp-TcaATcaB from that of the other TcAs (Supplementary Figure 12c). Due to one loop per domain that is 5-8 residues shorter in Xn-XptA1, Mm-TcdA4, and Yp-TcaATcaB than in Pl-TcdA1 (Supplementary Figure 12d,e), the shells of these proteins are not closed at their bottom (Figure 6a-c). Therefore, this region cannot be responsible for holding together the shell domains in the prepore pentamer.

The surface electrostatic potentials at pH 4 and pH 11 of the neuraminidase-like domain of all TcAs show an accumulation of either positive or negative charges leading to repulsions at these pH values (Figure 6a,c). At neutral pH, however, only Yp-TcaATcaB shares the equal distribution of opposite charges with Pl-TcdA1 (Figure 6b). Xn-XptA1 and Mm-TcdA4 are more negatively or positively charged in this region, respectively (Figure 6b), indicating that the shell is not stabilized by complimentarily charged residues in the neuraminidase-like domains. Nevertheless, these TcAs formed stable prepore complexes at neutral pH and the prepore-to-pore transition could be induced by changing the pH (Supplementary Figure 13).

Thus, due to the differences in charge distribution among TcA neuraminidase-like domains as well as the relatively wide opening at the bottom of some of the TcA shells, we conclude that this domain is not responsible for the electrostatic lock mechanism that has been observed in all studied TcAs (Supplementary Figure 13).

We therefore analyzed the interfaces between two shell domains in order to identify the conserved regions that could qualify as potential pH-sensitive elements. pH-dependent conformational changes have been shown to either implicate histidine-cation or anion-anion (glutamates and/or aspartates) interactions^26^. The involved residue pairs switch from stabilizing to destabilizing a conformation through pH-shift-induced electrostatic repulsion.

We could not find any conserved anion-anion pair or histidine-cation pair in the examined TcA structures. We only identified a prominent cluster of three histidines in Pl-TcdA1 (His 46 and 50 in protomer A and His 1808 in protomer B) (Supplementary Figure 14a,b). However, the residues are not conserved among the TcAs and the loop of protomer B containing H1808 is much shorter in Mm-TcdA4 and therefore out of range for an interaction with protomer A (Supplementary Figure 14c). As a result, we exclude that this histidine cluster is important for the function of the TcAs in general.

Since we could not find a classical pH-switch, we searched for other conserved ionic interactions in the shell and identified three buried pairs that are found at the interface between two protomers in all examined TcAs: E158 and R1873, D965 and R1971, and E1086 and R1166 (residue numbering of Pl-TcdA1) (Figure 6d-f). The interacting residues are in close spatial proximity in the prepore state (3.8 Å, 3.5 Å and 3.9 Å, respectively) and are likely forming salt bridges. In the Pl-TcdA1 pore state, none of these interactions are possible due to the enlarged distance after the conformational change (9.8 Å, 5.8 Å and 25 Å, respectively).

To analyze whether these interactions are involved in the pH-shift-induced destabilization of the shell, we designed three mutants of Pl-TcdA1 in which the respective residues are mutated to alanine (E158A-R1873A, D965A-R1971A and E1086A-R1166A). All mutants could be recombinantly expressed in *E. coli*. The purification of E158A-R1873A and D965A-R1971A yielded pure protein. Most of the E1086A-R1166A variant, however, aggregated during expression and we could only partially purify small amounts of the protein (Supplementary Figure 15a-c). For E158A-R1873A and D965A-R1971A, we observed the characteristic bell-shaped particles and the pore state could be induced by a pH shift to pH 11, indicating that these two variants behave like the wild type (Supplementary Figure 15a,b). Despite the impurity of Pl-TcdA1-E1086A-R1166A, we found TcAs in their pentameric prepore state in the sample (Supplementary Figure 15c). Interestingly, the pore state of the protein was also present (Supplementary Figure 15c), indicating that a part of the prepore complexes must have been transitioned to the pore state at neutral pH. This might also explain the aggregation during expression, since the pore state, where the hydrophobic transmembrane region is exposed, tends to aggregate in solution. Control mutants, where only one of the two residues was mutated to alanine, showed a similar expression pattern as the double mutant Pl-TcdA1-E1086A-R1166A (Supplementary Figure 16a). This suggests that the E1086-R1166 residue pair is crucial for the stability of the shell domain at neutral pH.

Based on these results, one might be tempted to speculate that this residue pair acts as the electrostatic lock of TcA that opens at low and high pH, thereby destabilizing the shell. However, the pKa of the side chain groups of glutamate is 4.25 and that of arginine is 12.48. The pKa values are even further shifted for residues involved in salt bridges ^27^. It is therefore unlikely that the residues change their protonation state at the pH values we used for inducing the pore state (pH 4.7 and pH 11, respectively).

Y1168, Y1205 and H1202 (numbering corresponding to Pl-TcdA1) are located in close vicinity to the E1086-R1166 residue pair (Supplementary Figure 16b). The three residues are conserved in Pl-TcdA1, Xn-XptA1, Mm-TcdA4, but not in Yp-TcaA-TcaB (Supplementary Figure 16c). To examine whether they have an influence on the interaction between E1086 and R1166, we mutated the tyrosines to phenylalanines (Pl-TcdA1-Y1168F-Y1205F) and the histidine to alanine (Pl-TcdA1-H1202A), and analyzed the TcA variants (Supplementary Figure 16d). We observed that the characteristic bell-shaped particles and the pore state could be induced by a pH shift to pH 11, indicating that these two variants behave like the wild type (Supplementary Figure 16d,e). Therefore, these residues are obviously not influencing the E1086-R1166 residue pair.

We conclude that TcAs, in general, do not have a classic pH switch. The conserved residue pair E1086-R1166 is essential for the stability of the complex at neutral pH. However, it cannot act as a pH switch due to its composition. We propose instead that the destabilization of many electrostatic interactions at the interface between the shells of two protomers are responsible for the opening of the shell at low and high pH values. The E1086-R1166 pair probably operates as a latch that springs open only once the other regions are destabilized by electrostatic repulsions.

## Conclusion

In our study, we investigated and compared five TcAs from four different organisms; *P. luminescens*, *X. nematophila*, *M. morganii* and *Y. pseudotuberculosis*. The cryo-EM structures of the five TcAs combined with mutational and functional studies revealed that their overall architecture and mechanism of action is similar. Important structural features relevant for the mechanism of action, i.e. the linker, the toxin translocation channel and the TcB-binding domain are present in all analyzed structures. These parts are the most conserved, both with respect to structural organization and residue conservation. All TcAs have a 3_1_ trefoil knot, that is strengthening the basis of the linker, which is especially put under strain during the prepore-to-pore transition. Therefore, the protein knot in TcA seems to be an important conserved structural feature of Tc toxins. In contrast, the β-sheet domains of the TcA shell, comprising the receptor-binding domains, are less conserved, parts of them are missing in some TcAs and show a higher structural flexibility than the rest of the protein.

Importantly, the TcA from the opportunistic human pathogenic bacterium *M. morganii*, which can cause various infections, such as abscess, sepsis, and nosocomial infections following surgery, resulting in a high mortality rate^28,29^ and the TcA from *Yersinia pestis*, the causative agent of plague^30^, are both functional Tc toxin components, suggesting that Tc toxins of these bacteria contribute as active toxins to the pathogenic effect. The structural difference at their periphery, i.e. the receptor-binding domains, indicates that these toxins adapted to the interaction with different host cells during their evolution, resulting in the host specificity observed for TcAs from different species^5^. In the extreme case of TcA from *Yersinia pestis*, which does not contain any receptor-binding domain, an additional coiled coil domain at the top of the shell compensates for the likely reduced stability of the complex due to the absence of receptor-binding domains.

TcAs are not only present in insects and human pathogenic bacteria, but they have been also identified in plant pathogenic bacteria such as *Pseudomonas syringae* and *Pseudomonas fluorescens*^31^. Although we have not studied TcA from these bacteria, based on our study, we propose that all TcAs, including those of plant pathogenic bacteria share the same architecture and mechanism of action. The high structural conservation of Tc toxins allows the formation of functional chimeric holotoxins, opening up new avenues for the design of potential biopesticides.

An important step in the mechanism of action of Tc toxins is the destabilization of a pH-sensitive electrostatic lock in the shell that releases the channel at high or low pH values^11^. Since we could not identify a classic pH switch, we propose that many electrostatic interactions at the shell-shell interfaces are responsible for the opening of the shell at low and high pH values instead. A conserved ionic pair that stabilizes the shell, likely operates as a strong latch that only springs open after the electrostatic destabilization of other regions.

Taken together, our study provides a holistic view on the architectural organization and function of Tc toxins that leads to new insights into the architecture and host specificity of the Tc toxin family.

## Acknowledgements

We thank A. Elsner for purification of Mm-TcdA4 and Pl-TcdA1 and for assistance with site-directed mutagenesis. We thank K. Vogel-Bachmayr for the purification of Pl-TcB-TcC. We acknowledge the help of F. Merino during analysis of ionic interaction studies and B. Rath for the preliminary structure investigations of Yp-TcaATcaB. We gratefully thank O. Hofnagel and D. Prumbaum for their excellent assistance in electron microscopy. We gratefully acknowledge R. Matadeen and S. de Carlo (FEI Company) for image acquisition of the Pl-TcdA4 data set at the Netherlands Centre for Nanoscopy in Leiden (NeCEN). This work was supported by funds from the Max Planck Society (to S.R.) and the European Research Council under the European Union’s Seventh Framework Programme (FP7/2007-2013) (grant no. 615984) (to S.R.).

## Author Contributions

S. R. designed and supervised the project. F.L. processed and analyzed cryo-EM data and built atomic models of Xn-XptA1, Mm-TcdA4 and Yp-TcaATcaB. C.G. and D.R. processed Pl-TcdA4 and Pl-TcdA1 data, respectively, and D.R. built the atomic model of Pl-TcdA1. F.L. designed the proteins, performed structural analysis and mutational studies. R.B. and F.L. performed single-channel conductivity experiments. D.R. performed BLI and intoxication experiments. D.M. analyzed the organization of the α-helical shell. F.L. prepared figures, F.L. and S.R. wrote the manuscript.

## Author Information

The cryo-EM densities of Pl-TcdA1, Pl-TcdA4, Xn-XptA1, Mm-TcdA4 and Yp-TcaATcaB have been deposited in the Electron Microscopy Data Bank under accession numbers … and …, respectively. The corresponding coordinates have been deposited in the Protein Data Bank under accession number…and …, respectively. Correspondence and requests for materials should be addressed to S.R.

## Supplementary Figures

**Supplementary Figure 1:**
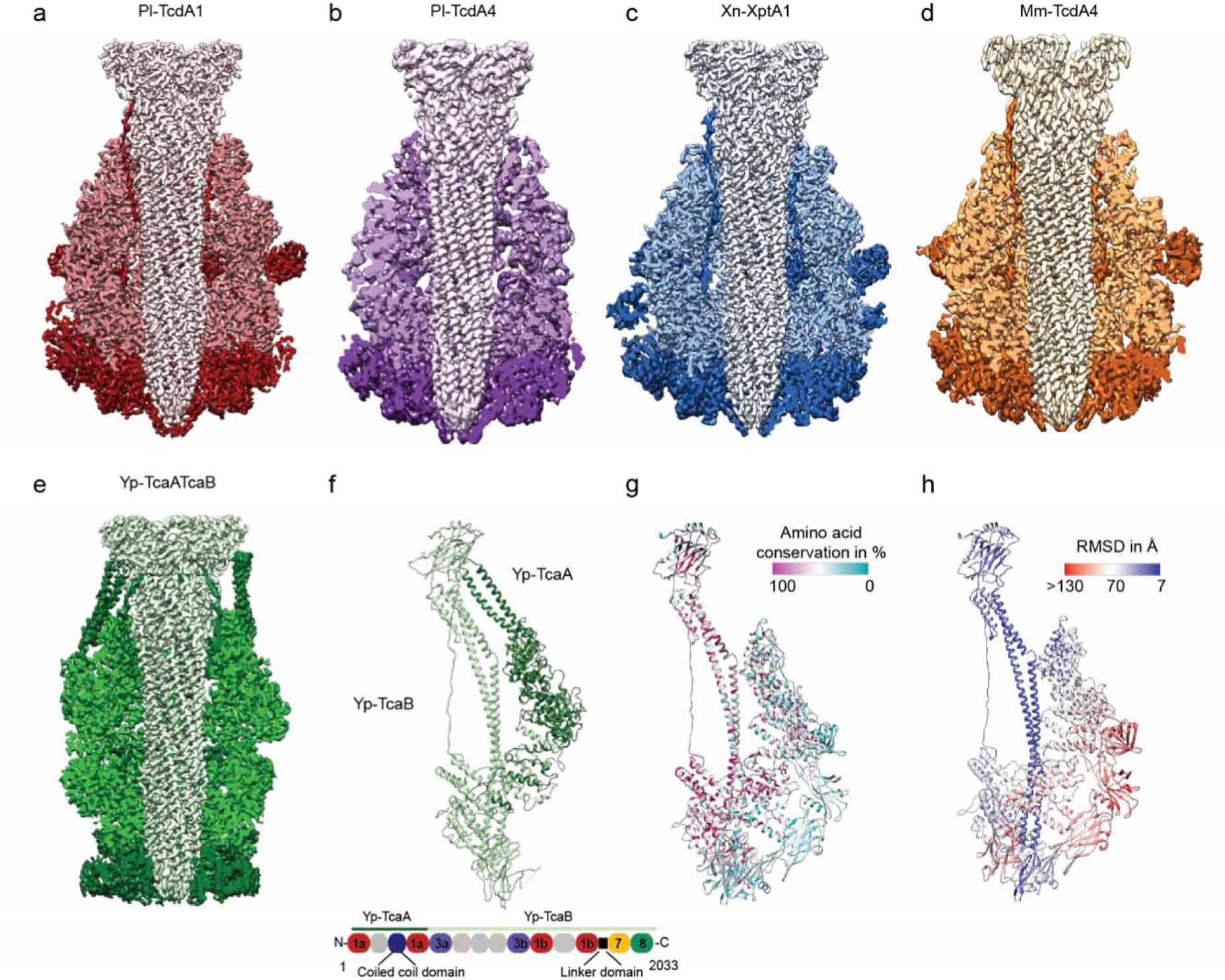
Structure and conservation of TcAs. (**a-e**) Longitudinal section through the density maps of the five TcAs to display the pore-forming channel. The outer shell encloses the tip of the channel completely in Pl-TcdA1 and Pl-TcdA4, but not in Xn-XptA1, Mm-TcdA4 and Yp-TcaATcaB. Color gradient from light to dark according to Figure 1. (**f**) Structure of a Yp-TcaATcaB protomer showing the two components Yp-TcaA (light green) and Yp-TcaB (dark green) as well as the domain sequence. (**g**) Structure of a Pl-TcdA1 protomer demonstrating the conservation of residues between Pl-TcdA1, Xn-XptA1, Pl-TcdA4, Mm-TcdA4 and Yp-TcaATcaB. Highly conserved residues are depicted in magenta, less conserved residues are depicted in cyan. (**h**) Structure of a Pl-TcdA1 protomer colored according to the root mean square deviation (RMSD) values between the five TcAs. Regions with high and low RMDs are depicted in red and blue, respectively. See also Supplementary Video 1.

**Supplementary Figure 2:**
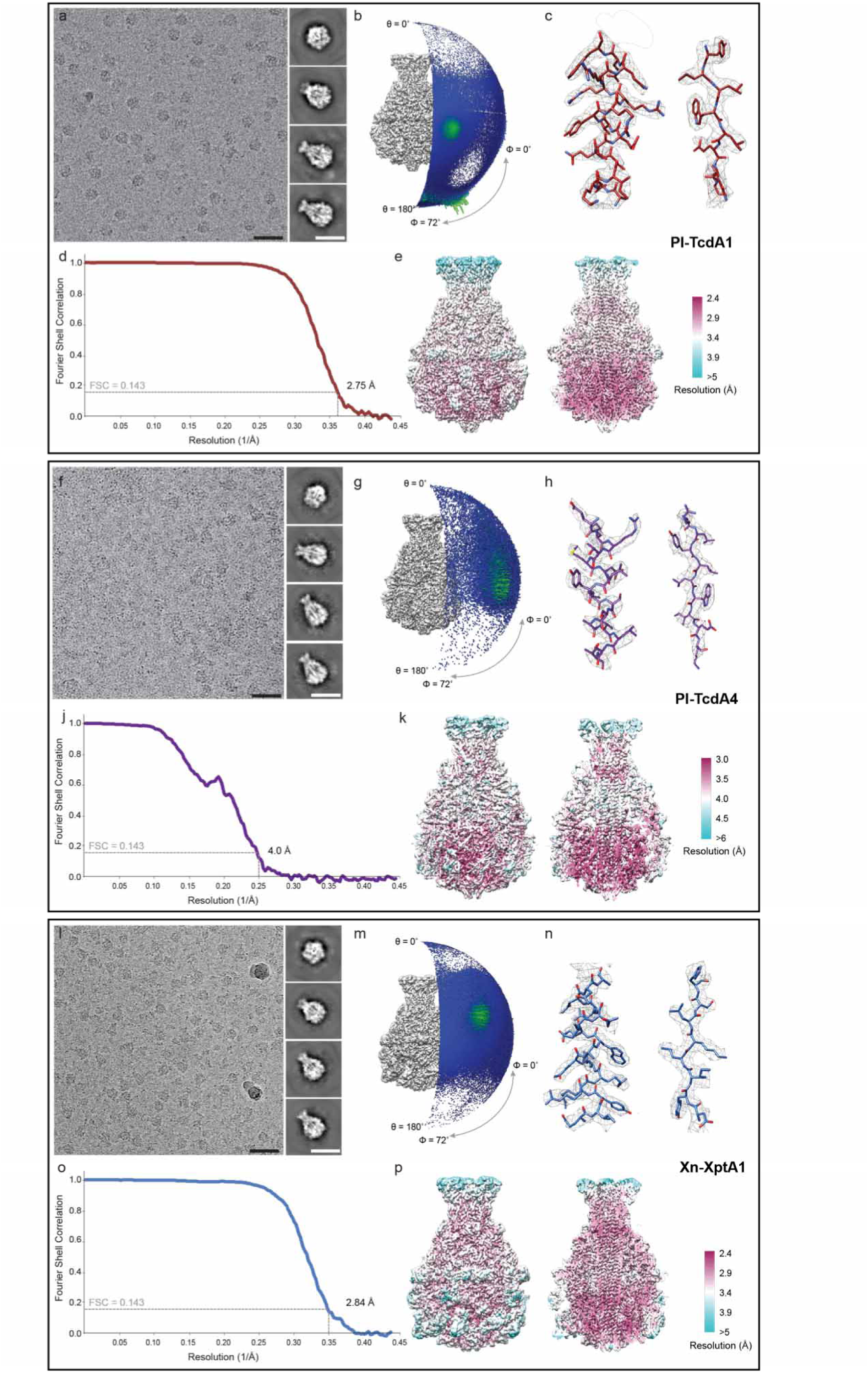
EM analysis of Pl-TcdA1, Pl-TcdA4, and Xn-XptA1. (a, f, l) Typical motion-corrected micrographs (scale bars, 50 nm) and 2D class averages (scale bars, 25 nm). (b, g, m) Angular distribution of all particles used for the final reconstruction. (c, h, b) Cryo-EM density (mesh) with the fitted atomic model, showing an α-helical part (left) and a β-strand region (right). (d, j, o) Fourier shell correlation (FSC) curves of the final, filtered density maps. The average resolution at 0.143 FSC criterion is indicated. (e, k, p) EM density maps colored according to the local resolution, showing the complete electron density and a longitudinal cut through the density maps.

**Supplementary Figure 3:**
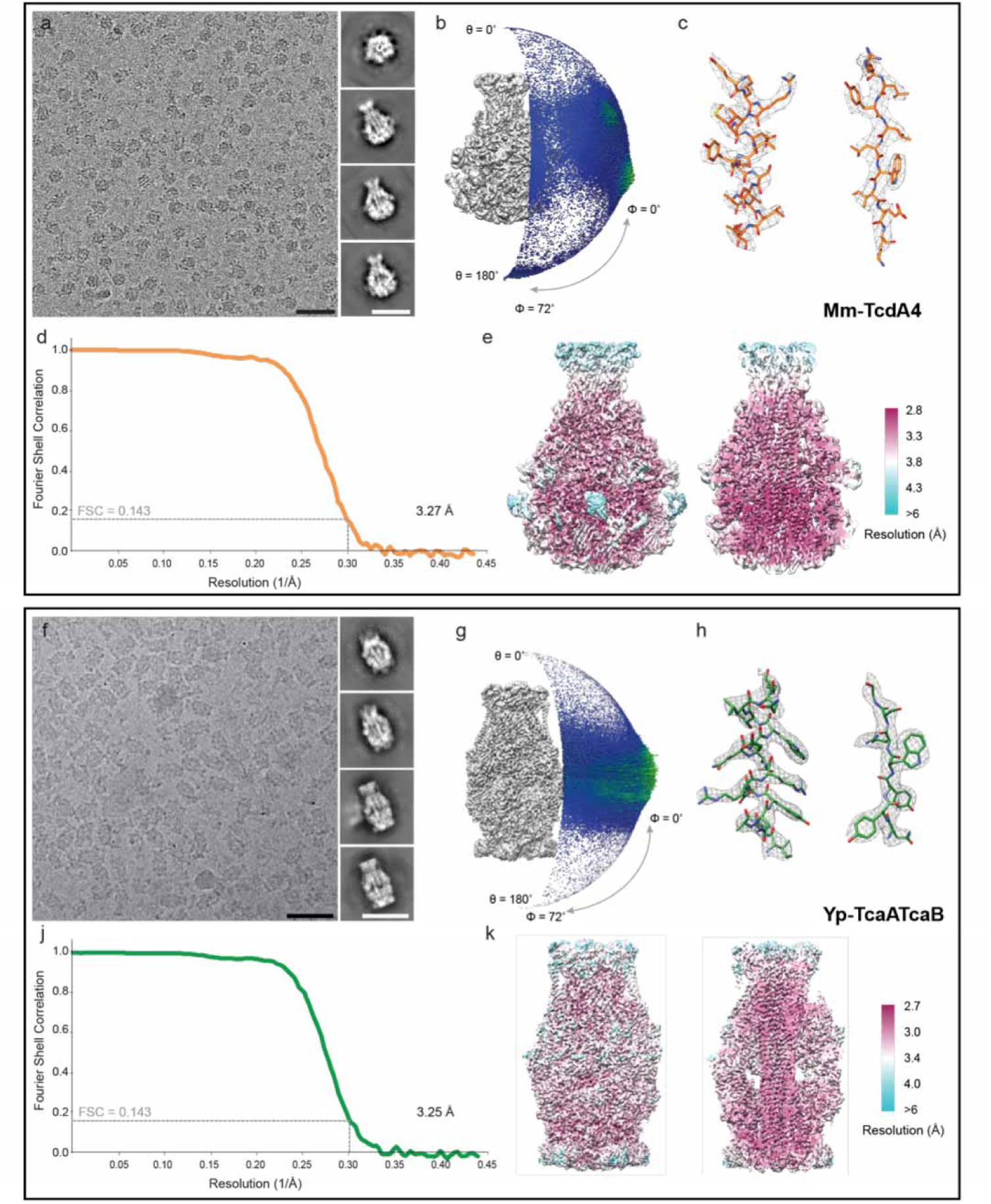
EM analysis of Mm-TcdA4 and Yp-TcaATcaB. (a, f) Typical motion-corrected micrographs (scale bars, 50 nm) and 2D class averages (scale bars, 25 nm). (b, g) Angular distribution of all particles used for the final reconstruction. (c, h) Cryo-EM density (mesh) with the fitted atomic model, showing an α-helical part (left) and a β-strand region (right). (d, j) Fourier shell correlation (FSC) curves of the final, filtered density maps. The average resolution at 0.143 FSC criterion is indicated. (e, k) EM density maps colored according to the local resolution, showing the complete electron density and a longitudinal cut through the density maps.

**Supplementary Figure 4:**
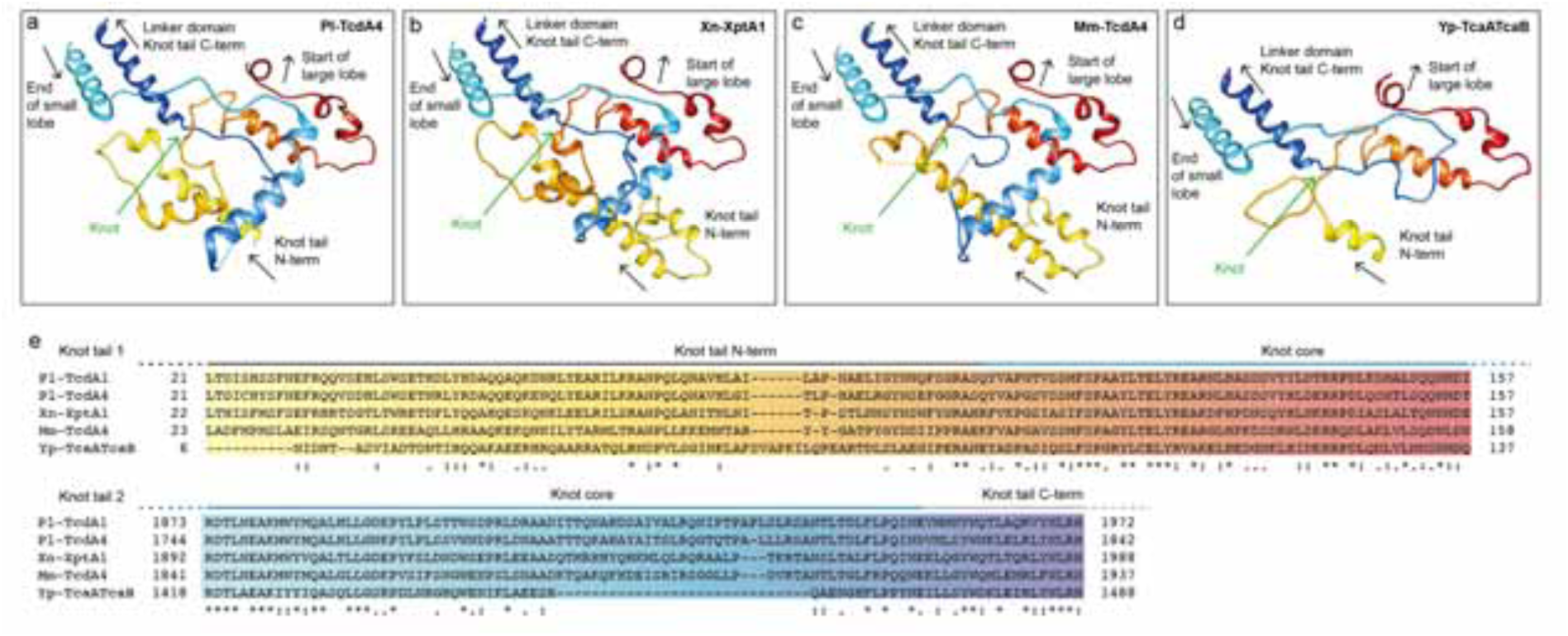
A trefoil 31 protein knot is present in all five TcAs. (a-d) Close-up view of the protein knot of Xn-XptA1 (a), Pl-TcdA4 (b), Mm-TcdA4 (c) and Yp-TcaATcaB (d), respectively. All protein knot structures are depicted in a color gradient. The arrows indicate the direction of sequence. The C-terminal knot tail is colored in a red-to-yellow gradient and the N-terminal knot tail is colored in a dark-to-light blue gradient. (e) Protein sequence alignment of the two tails of the protein knot. The color gradient is identical to panels a-d and the location of the knot regions are depicted in dark grey, blue and light grey, respectively. See also Supplementary Figure 5 and Supplementary Video 2.

**Supplementary Figure 5:**
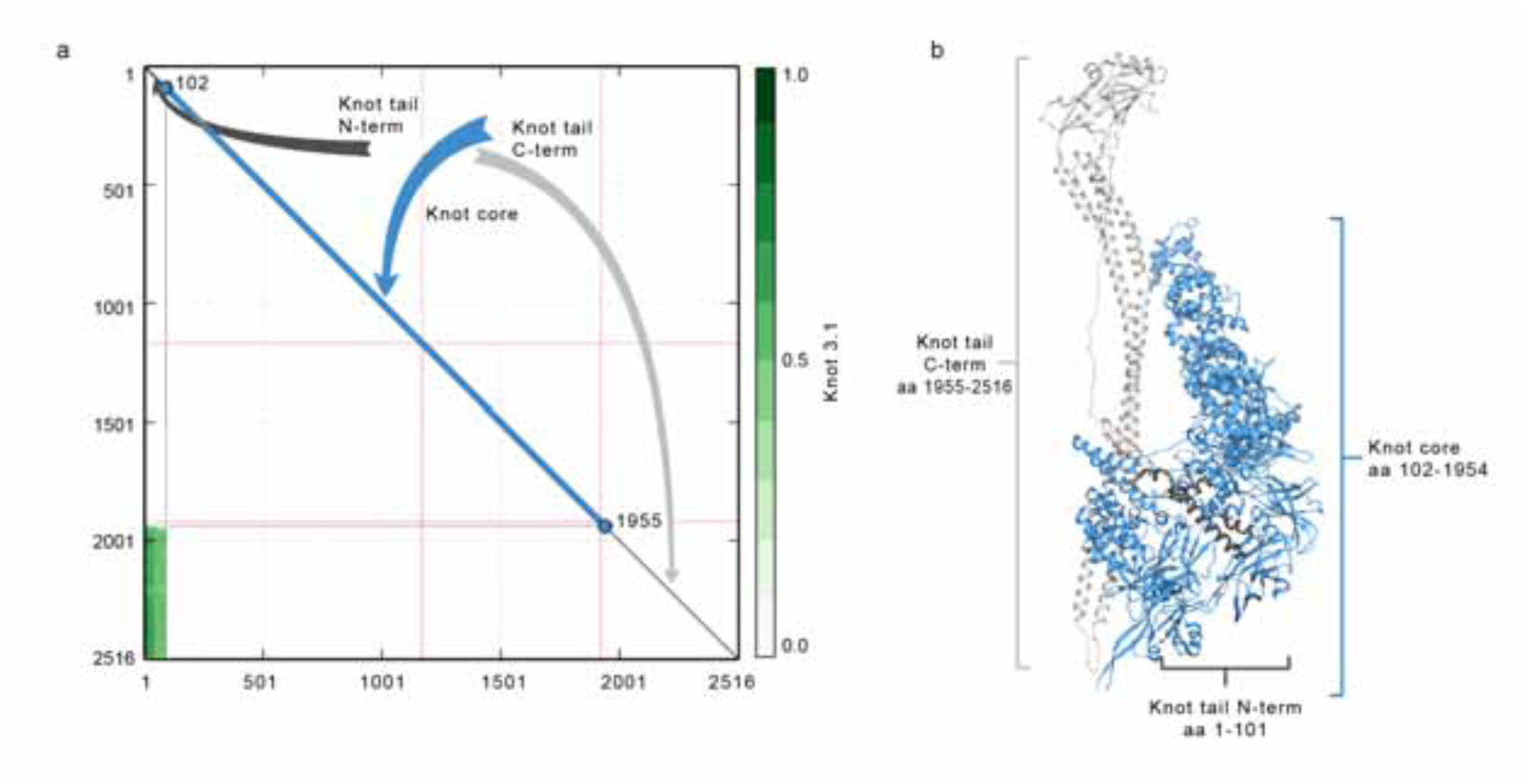
Organization of the trefoil 31 protein knot for Pl-TcdA1. (a) Matrix scheme of the trefoil 31 protein knot with the three domains of the knot organization: knot tail N-term (dark grey), the knot core (blue) and the knot tail C-term (grey). Residues belonging to the knotted chain are colored green. (b) Structure of a Pl-TcdA1 protomer colored according to the three different knot domains.

**Supplementary Figure 6:**
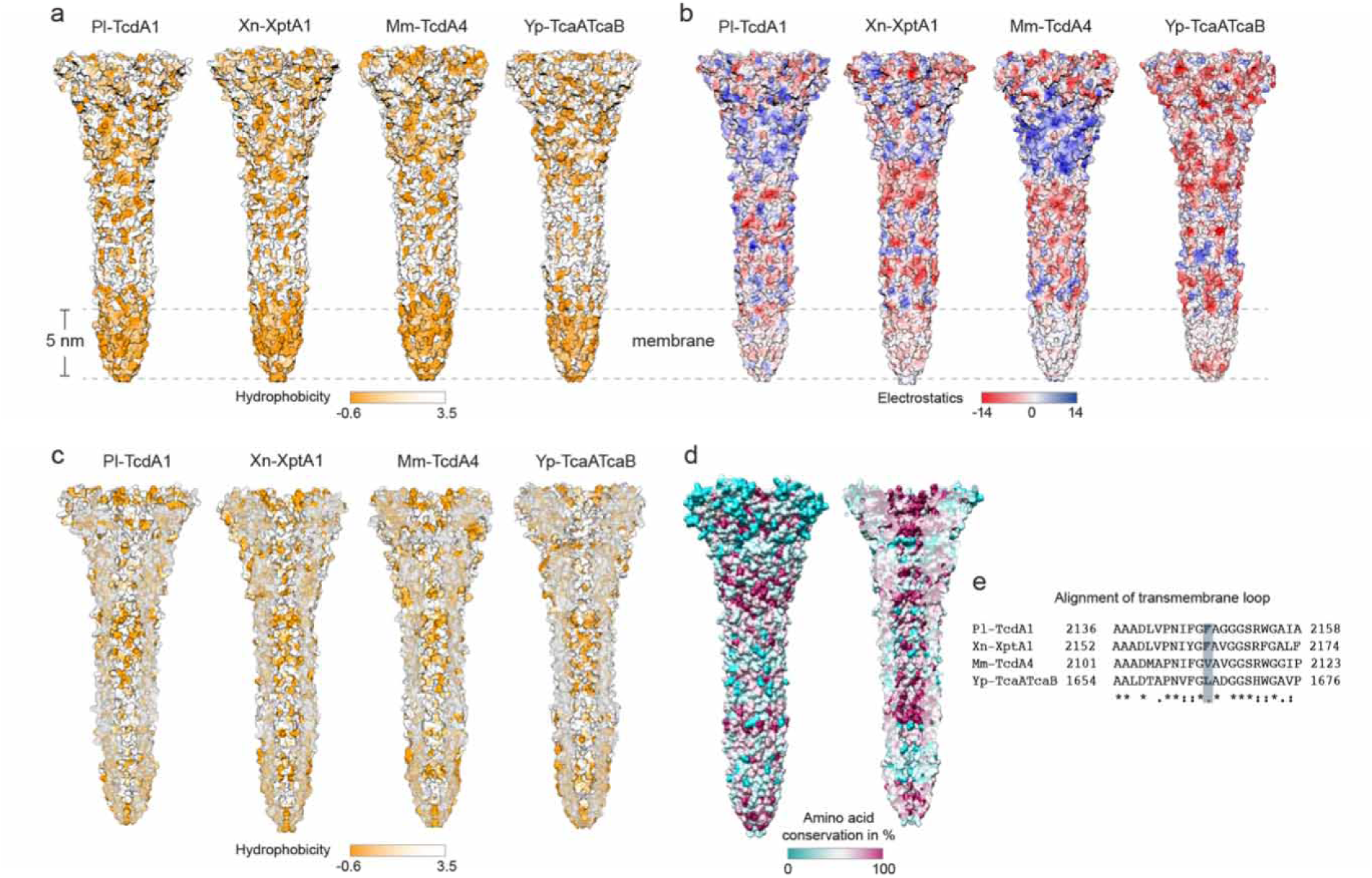
Biophysical properties of the TcA channel (a) Surface representation of the channels of Pl-TcdA1, Xn-XptA1, Mm-TcdA4 and Yp-TcaATcaB, colored by hydrophobicity. (b) Surface representation of the channels showing the electrostatic Coulomb potential at pH 7. Positively charged (14 kcal/mol) and negatively charged (−14 kcal/mol) residues are colored in blue and red, respectively. (c) Cross-sections of the channels showing the hydrophobicity at the channel inner surface. (d) Residue conservation plot of the channel domains of the four TcAs, shown for the channel surface (left) and the lumen of the channel (right). Conserved residues are shown in magenta. (e) Sequence alignment of the transmembrane loop of the channel domains. The residue at the very tip of the channel domain is highlighted in light blue.

**Supplementary Figure 7:**
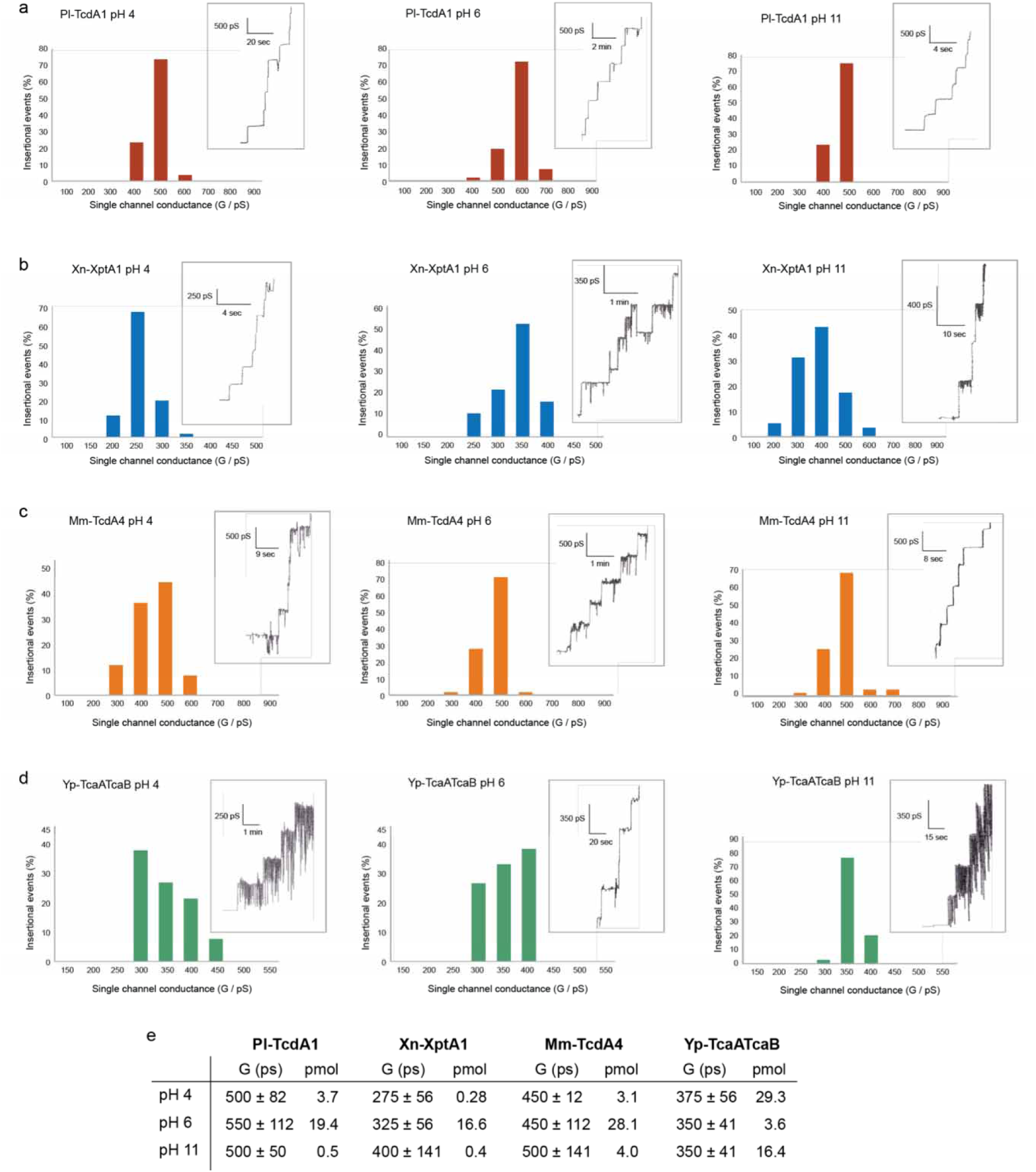
Distribution of single channel conductance of TcAs at different pH values. All histograms were constructed from the data of at least 70 pore insertional events for each TcA and pH value. (a-d) Measurements of Pl-TcdA1, Xn-XptA1, Mm-TcdA4 and Yp-TcaATcaB were performed at pH 4, 6 and 11, respectively. For each experiment, an exemplary recording of the single channel current versus time is attached. (e) Table listing the average single channel conductivity and the amount of TcA used for each experiment to obtain at least 70 pores.

**Supplementary Figure 8:**
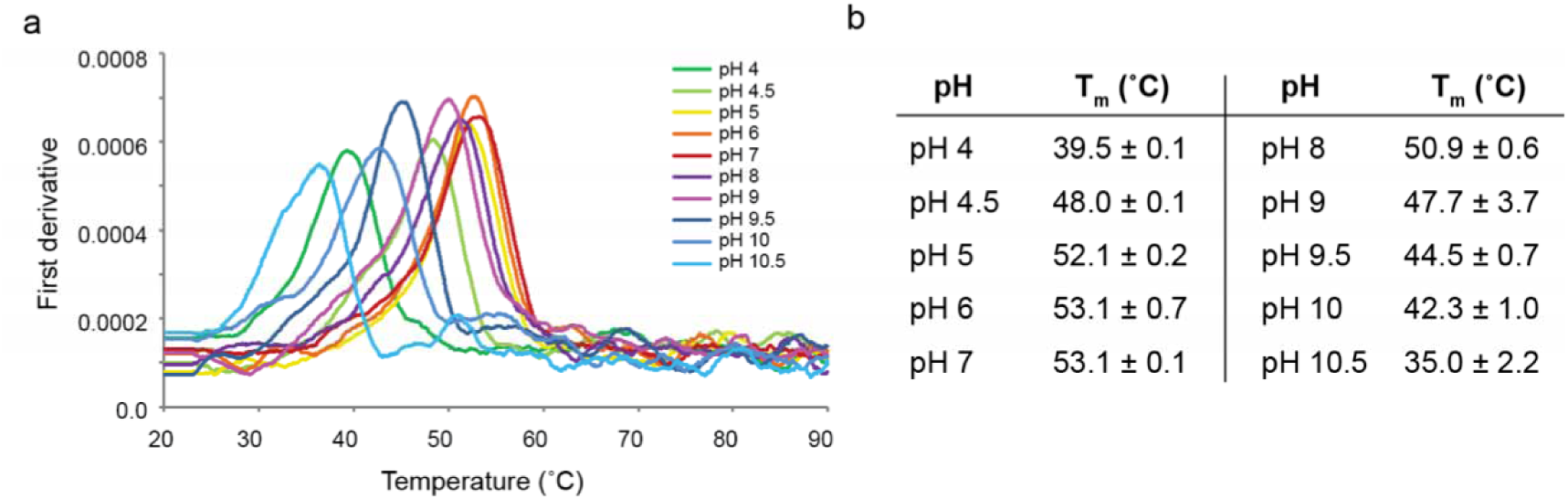
Nano differential scanning fluorimetry of Yp-TcaATcaB at different pH values. (a) The graph shows the first derivate of the absorbance quotient (330 nm/350 nm) as function of temperature. The different pH values are indicated by different colors. (b) Table with the melting temperatures (Tm) for each pH value. The measurements were performed in triplicates and the Tm is given as the mean ± standard deviation.

**Supplementary Figure 9:**
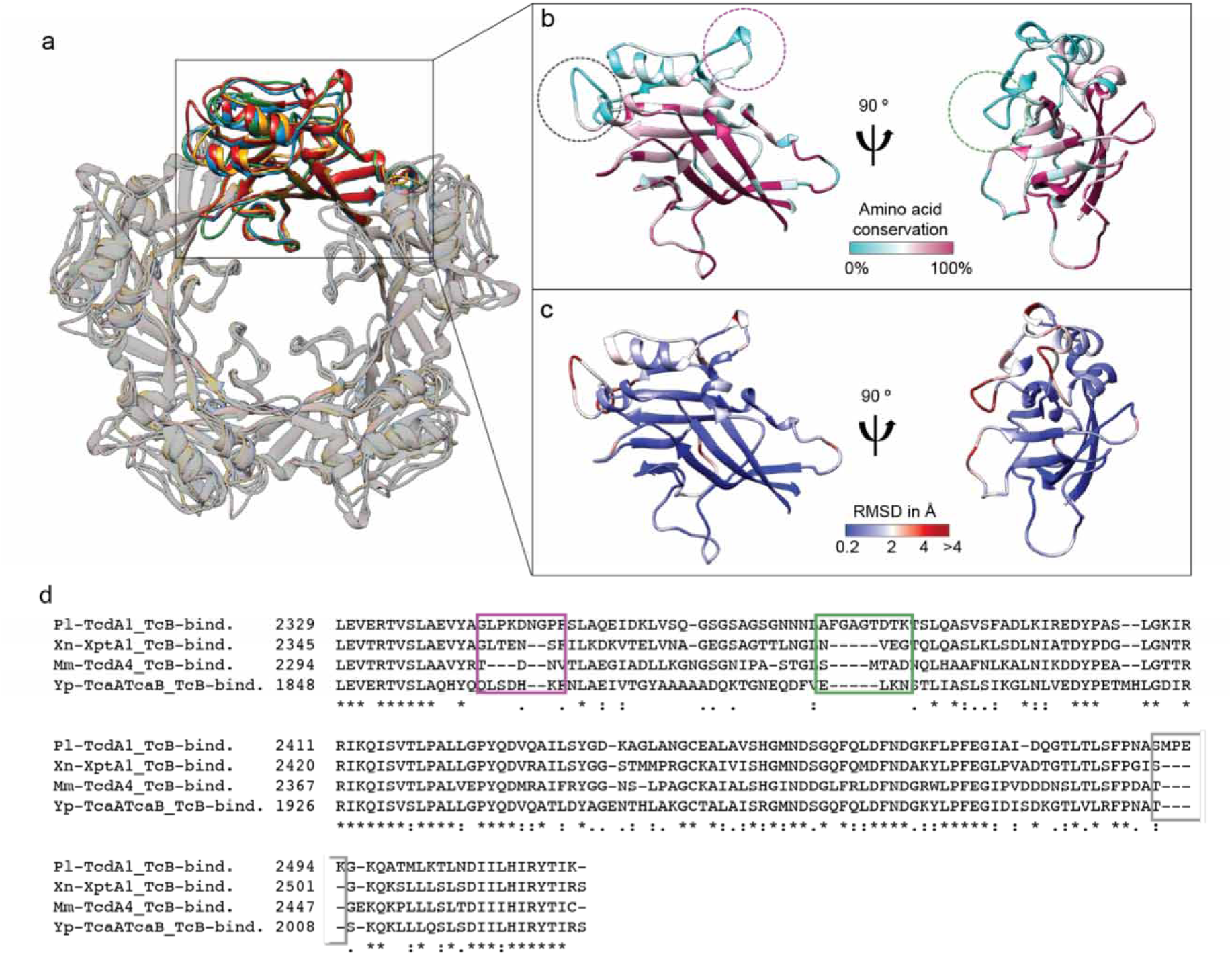
Comparison of the TcB-binding domains. (a) Overlay of the TcB-binding domains of Pl-TcdA1, Xn-XptA1, Mm-TcdA4 and Yp-TcaATcaB colored in red, blue, orange and green, respectively, in the context of the pentamer. One protomer is highlighted. (b) TcB-binding domain of one protomer of Pl-TcdA1 demonstrating the conservation of residues between Pl-TcdA1, Xn-XptA1, Mm-TcdA4 and Yp-TcaATcaB. Conserved residues are depicted in magenta. The three non-conserved loops are highlighted with dashed circles and the respective sequences are highlighted in (d). (c) RMSD of the TcB-binding domain structures. Residues with a low RMSD are colored in blue. (d) Sequence alignment of the TcB-binding domain. The loops marked in (b) are indicated in the alignment.

**Supplementary Figure 10:**
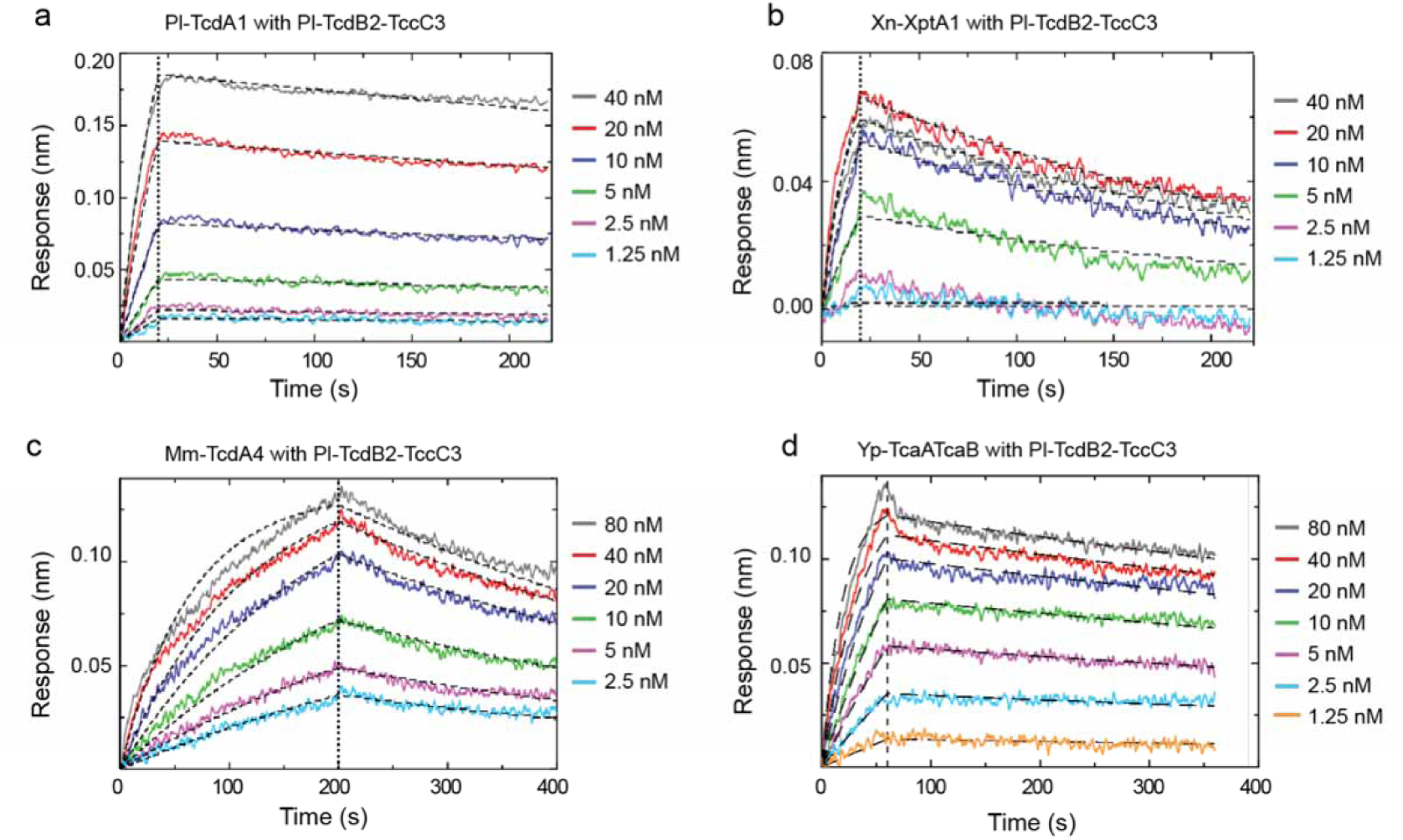
Binding affinities of chimeric holotoxins. (a-d) BLI sensorgrams of Pl-TcdA1 (a), Xn-XptA1 (b) Mm-TcdA4 (c) and Yp-TcaATcaB (d) interacting with immobilized Pl-TcdB2-TccC3. TcA pentamer concentrations were 1.25 – 40 nM in a and b, 2.5 – 80 nM in **c** and 1.25 – 80 nM in **d**. A global fit according to a 1:1 binding model was applied (black dashed curves). Association and dissociation phases are separated by a black dashed line. The obtained K_D_, k_on_ and k_off_ values are shown in Figure 5e.

**Supplementary Figure 11:**
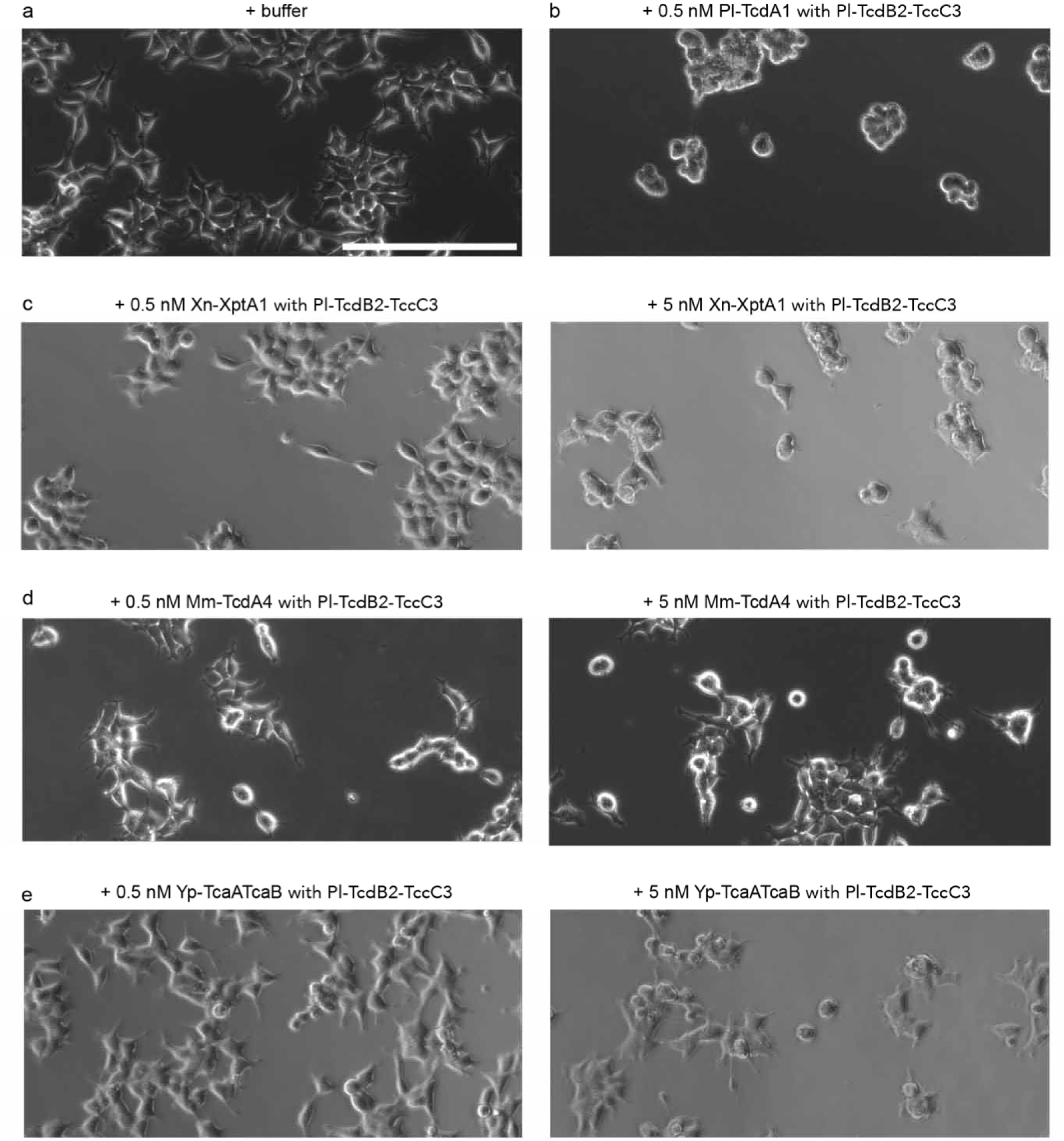
Intoxication of HEK293T cells with chimeric holotoxins. (a) Negative control with only buffer added. (b) Positive control with the native holotoxin complex from P. luminescens (Pl-TcdA1 and TcdB2-TccC3) at a concentration of 0.5 nM holotoxin. (c-**e**) Effect of the holotoxin complex composed of Pl-TcdB2-TccC3 and Xn-XptA1 (**c**), or Mm-TcdA4 (**d**), or Yp-TcaATcaB (**e**) at concentrations of 0.5 nM (left) and 5 nM (right). Cells were imaged 20 hours after infection. Scale bar, 200 µm.

**Supplementary Figure 12:**
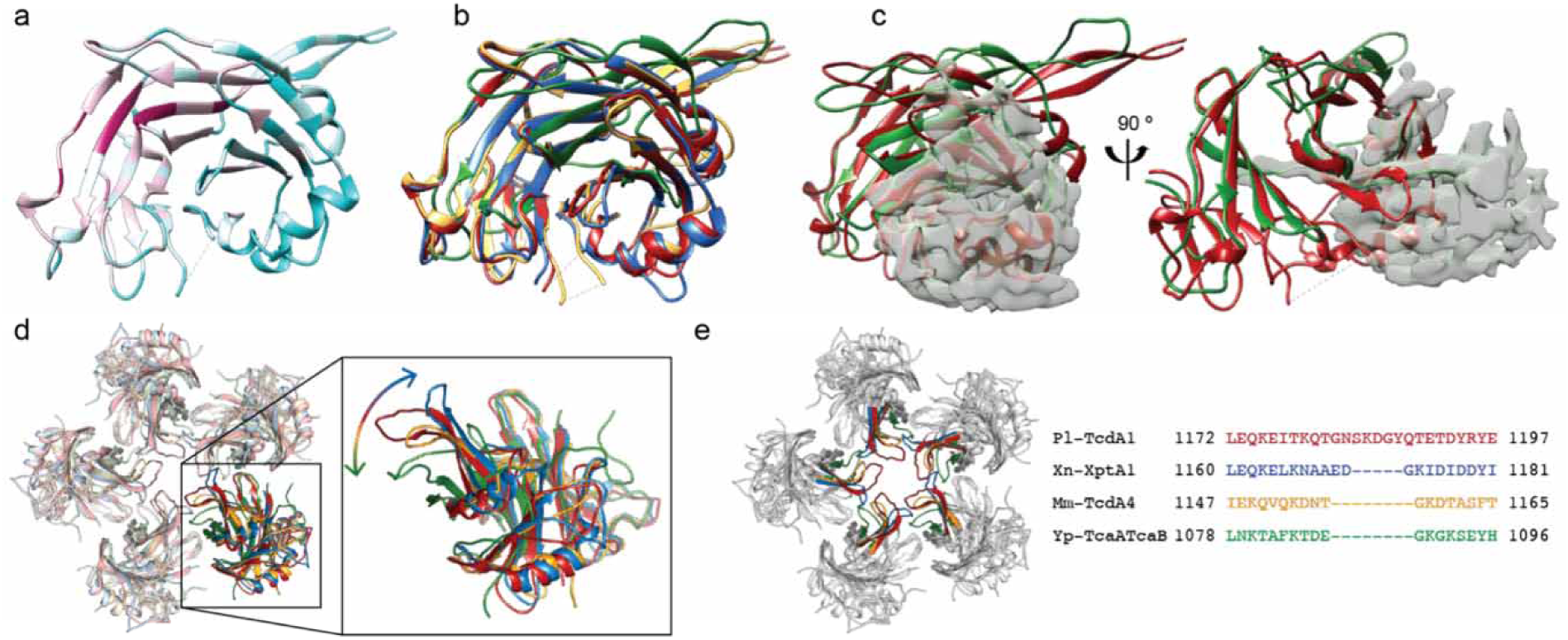
Topology of the neuraminidase-like domain. (a) Conservation of the neuraminidase-like domain with highly conserved regions in magenta and low conservation in turquoise. (b) Structural alignment of the domain for all four TcAs with respective colors (Pl-TcdA1 in red, Xn-XptA1 in blue, Mm-TcdA4 in yellow and Yp-TcaATcaB in green). (c) Structural alignment of the neuraminidase-like domain of Pl-TcdA1 and Yp-TcaATcaB shown in front (left) and side view (right). To visualize the missing 125 residues of Yp-TcaATcaB, the density map corresponding to these residues is shown. (d) Pentamer of the neuraminidase-like domain of Pl-TcdA1 (red), Xn-XptA1 (blue), Mm-TcdA4 (yellow) and Yp-TcaATcaB (green) with one protomer highlighted and zoomed in. The arrow indicates the different orientation of the flexible loop at the center of the pentamer. (e) Pentamer of the neuraminidase-like domain, highlighting the flexible loop and a sequence alignment showing the different lengths of the loop region in respective colors.

**Supplementary Figure 13:**
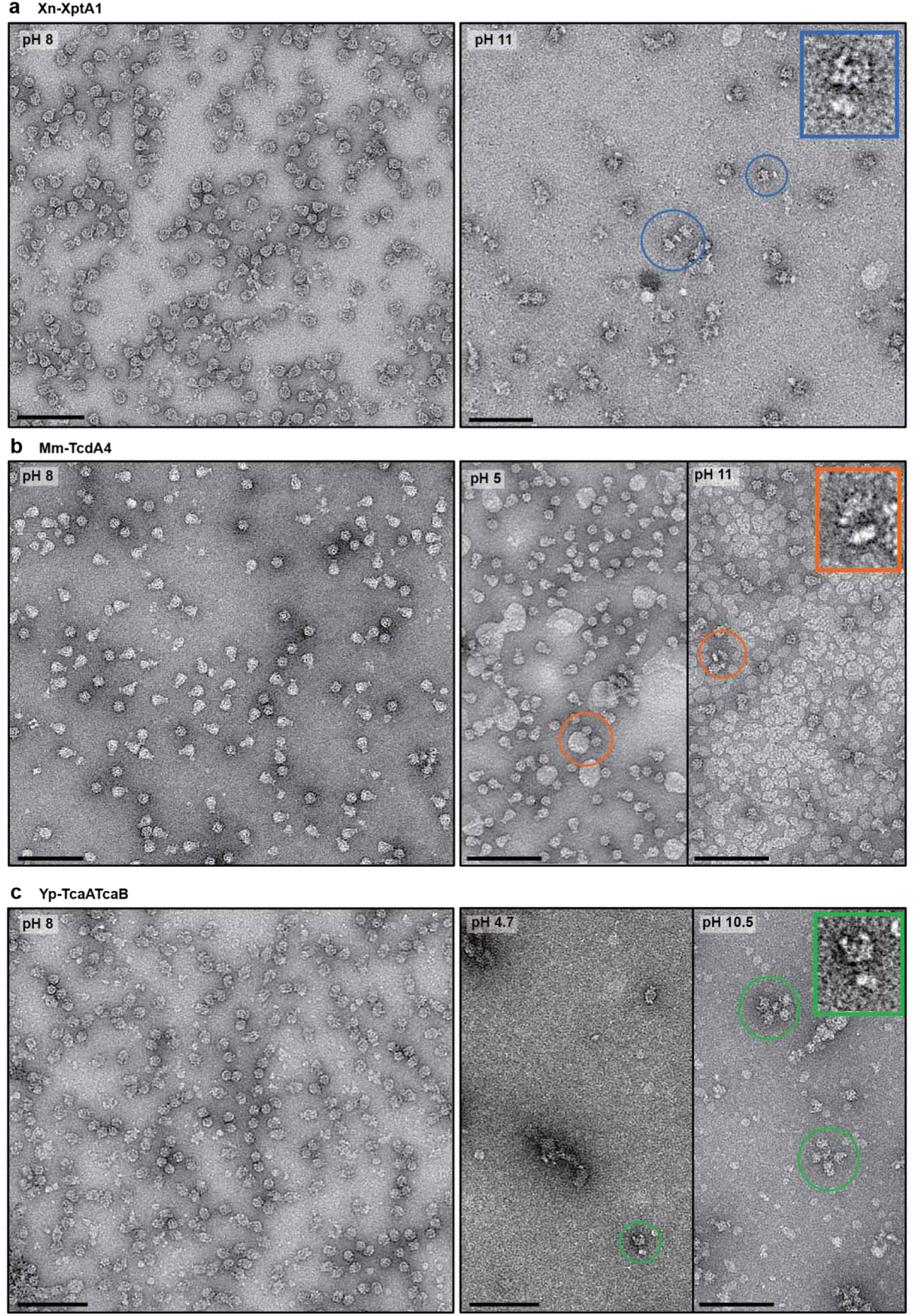
pH-induced pore formation of Xn-XptA1, Mm-TcdA4 and Yp-TcaATcaB. (a) Negative stain electron micrograph of the Xn-XptA1 prepore at pH 8 (left) and the pore reconstituted in nanodiscs at pH 11 in the presence of 3 mM CaCl2 (right). (b) Negative stain electron micrographs of the Mm-TcdA4 prepore at pH 8 (left) and the pore reconstituted in liposomes at pH 5 or in nanodisc at pH 11 and 5 mM CaCl2 (right). (c) Negative stain electron micrographs of the Yp-TcaATcaB prepore at pH 8 (left) and the pore reconstituted in nanodiscs at pH 4.7 or pH 10.5 (right). Particles in the pore state are highlighted with colored circles. One typical particle in the pore state is magnified for each protein.

**Supplementary Figure 14:**
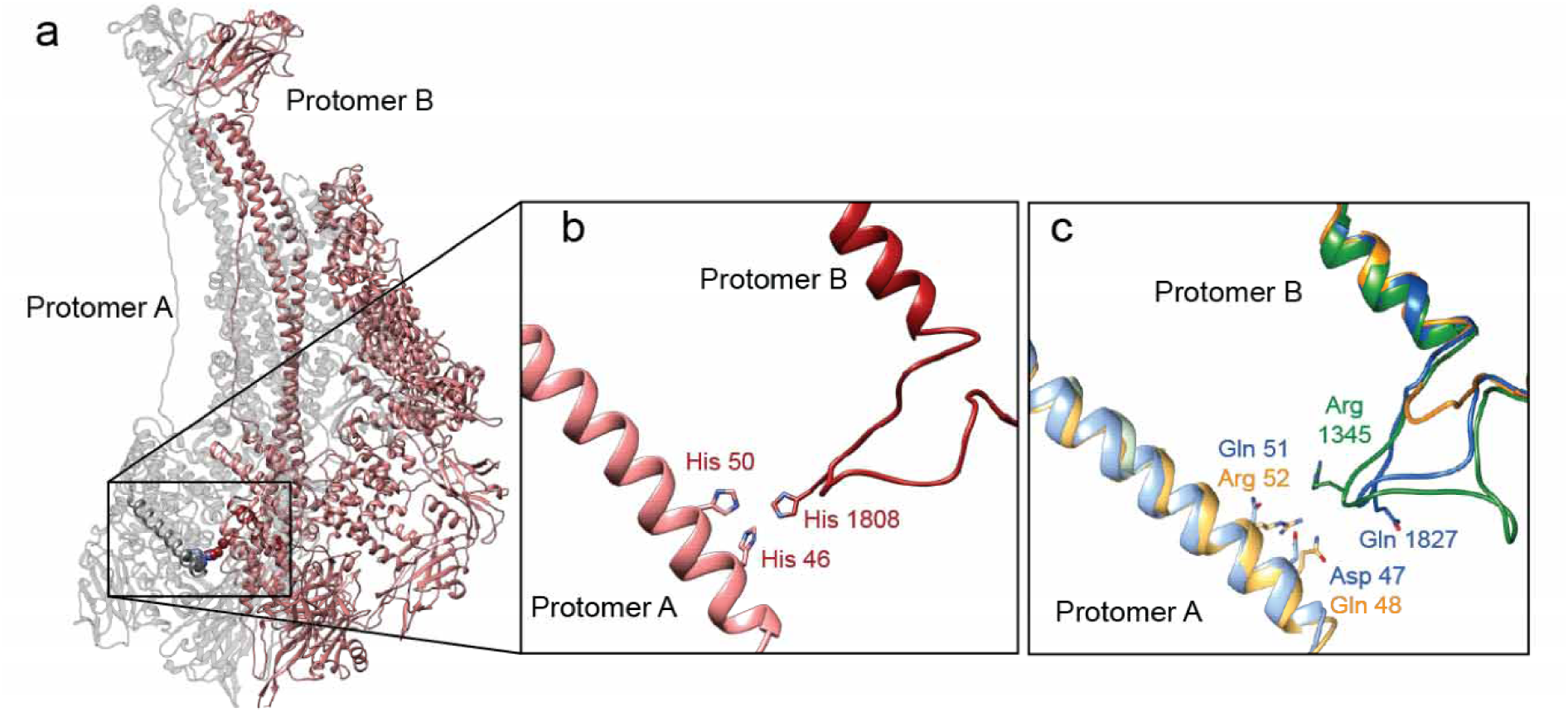
Non-conserved cluster of three histidine residues in Pl-TcdA1. (a) Two protomers of Pl-TcdA1 indicating the histidine interaction site between two protomers of Pl-TcdA1. (b) A close-up view of the three interacting histidines of Pl-TcdA1. (c) The histidine residues are not conserved in Xn-XptA1 (blue), Mm-TcdA4 (yellow) and Yp-TcaATcaB (green).

**Supplementary Figure 15:**
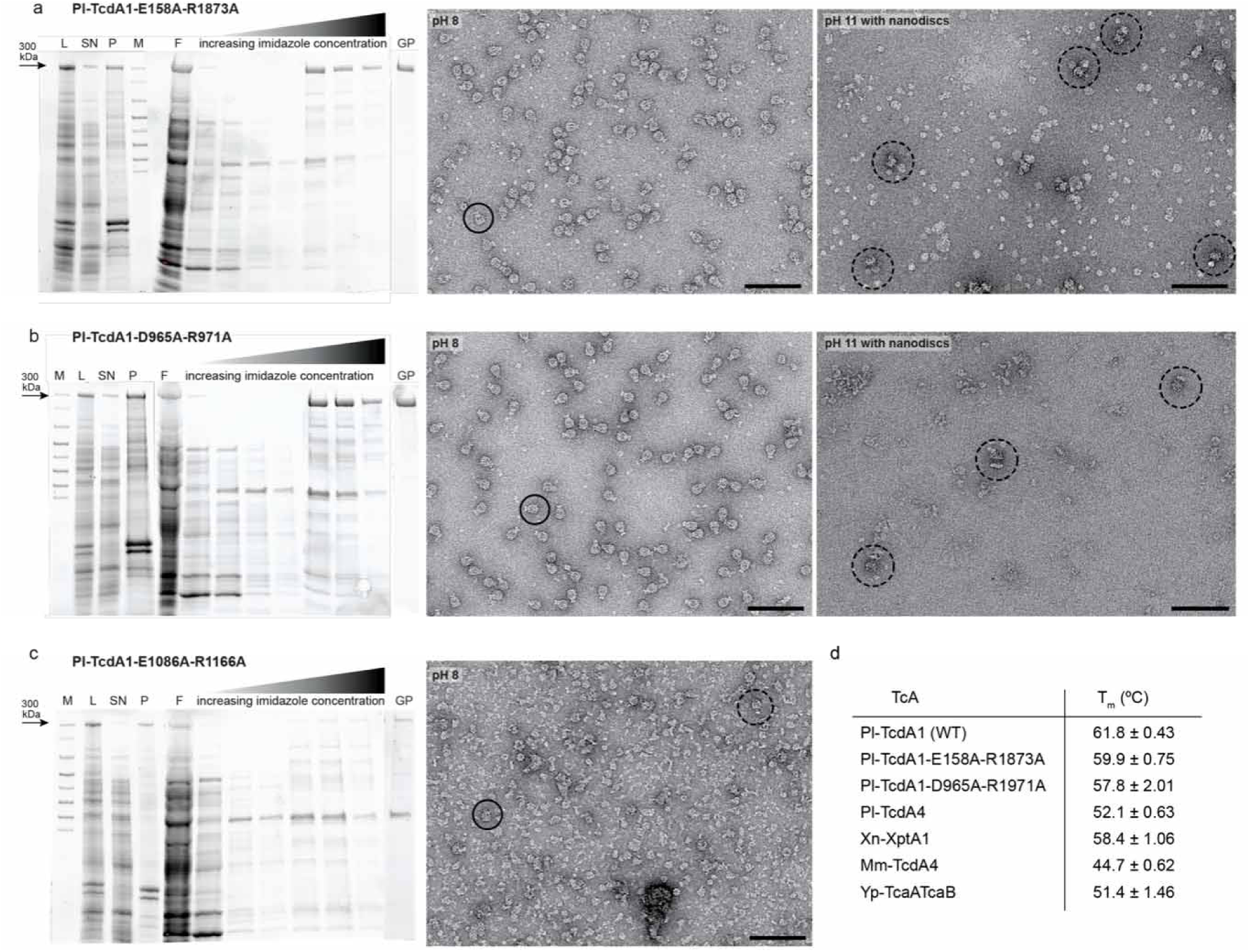
Alanine mutants of Pl-TcdA1 reveal differences in protein stability. (a-c) The left panels show an SDS-PAGE gel after Ni-NTA purification and gel filtration at pH 8 of Pl-TcdA1-D158A-R1873A (a), Pl-TcdA1D965A and R1971A (b), and Pl-TcdA1-D1086A-R1166A (c). M = marker, L = lysate, SN = supernatant, P = pellet, elutions at increasing imidazole concentrations ranging from 5 – 150 mM, GP = gel filtration peak fraction. The 300 kDa band corresponding to the TcA monomer is marked with an arrow. The right panels show negative stain electron micrographs of the peak fraction applied to the grid directly after purification at pH 8 or after reconstitution in nanodiscs at pH 11. Particles in the prepore and pore state are marked with solid and dashed circles, respectively. Due to the insufficient purity of the Pl-TcdA1-D158A-R1873A variant, the reconstitution step was omitted. (d) Table with measured melting temperatures Tm (nanoDSF) for all analyzed TcAs at pH 8. The measurements were performed as triplicates and the mean Tm as well as the standard deviation are shown.

**Supplementary Figure 16:**
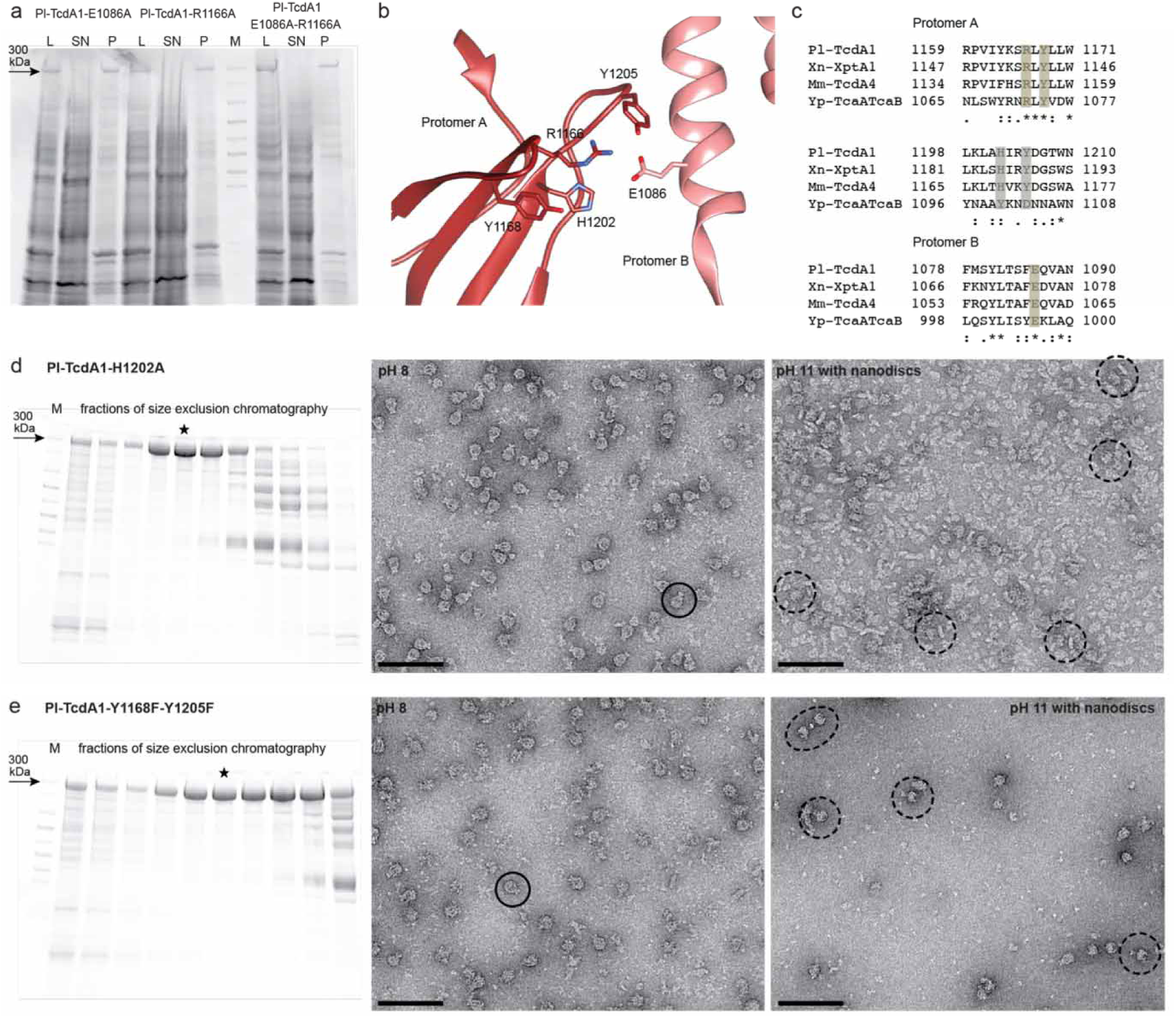
Mutational studies of Pl-TcdA1. (a) SDS-PAGE gel after expression of Pl-TcdA1 mutants Pl-TcdA1-E1086A, Pl-TcdA1-R1166A, and Pl-TcdA1-E1086A-R1166A. For all proteins, a sample of the lysate (L), supernatant (SN) and pellet after ultracentrifugation (P) was applied. M = marker. (b-c) Close-up view of a conserved interaction site in Pl-TcdA1 (b), and alignment of residues, which are involved in the interaction (c). The conserved residues in all four TcAs are highlighted in yellow and the ones only conserved in Pl-TcdA1, Xn-XptA1 and Mm-TcdA4 in green. (d-e) SDS-PAGE gel of gel filtration fractions (left) of Pl-TcdA1-H1202A (d) and Pl-TcdA1-Y1168F-Y1205F (e) as well as negatively stained complexes at pH 8 (middle) and pH 11 after reconstitution in nanodiscs (right). The ratio of toxin-nanodisc ratio was 1 : 10 in (d), leading to a background of empty nanodiscs. The star marks the band corresponding fraction used for electron microscopy. Particles in the prepore and pore state are marked with solid and dashed circles, respectively. Scale bars, 100 nm.

**Supplementary Table 1:**
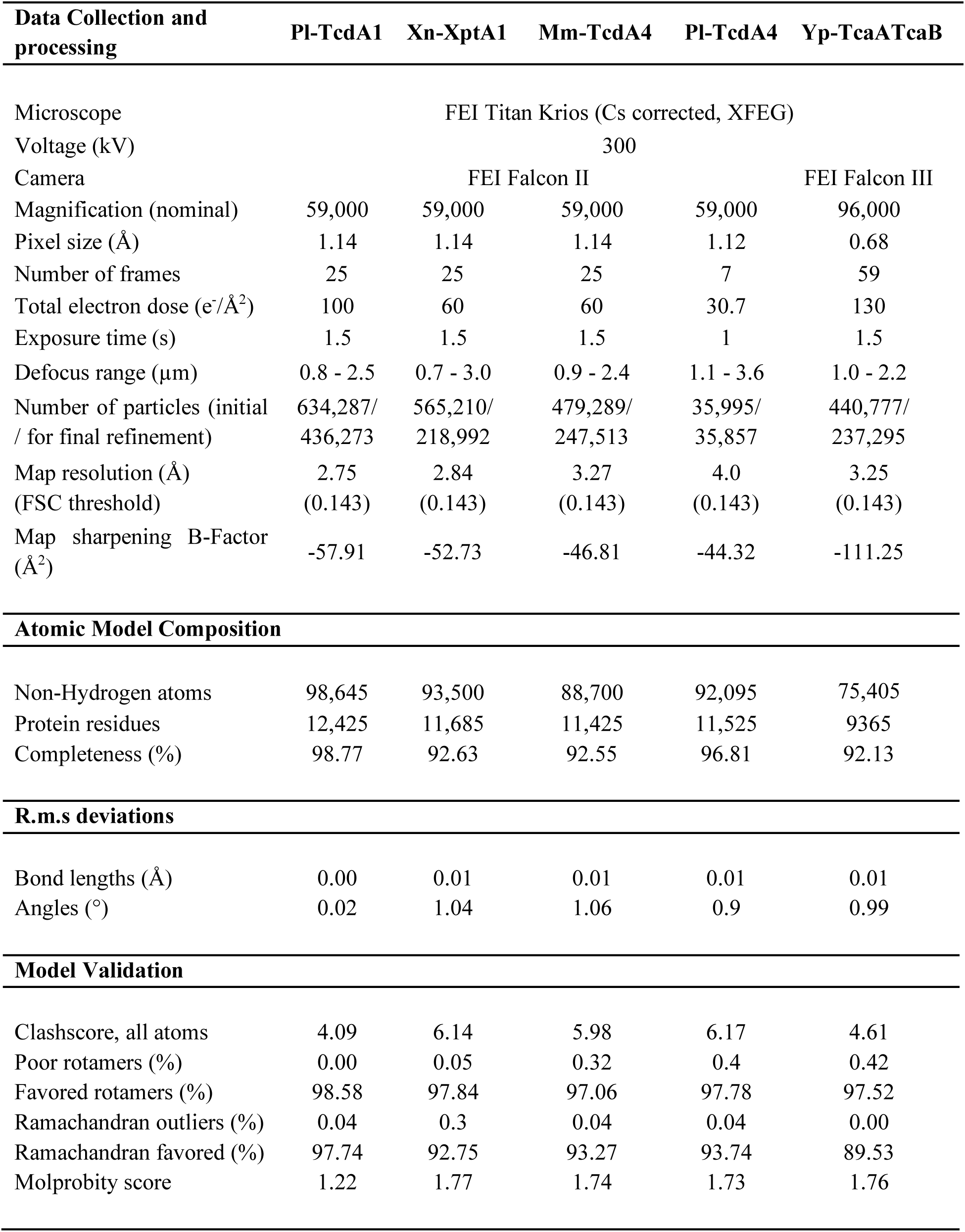
Statistics for cryo-EM data collection and model validation.

**Supplementary Video 1:** Cryo-EM density maps of Pl-TcdA1, Pl-TcdA4, Xn-XptA1, Mm-TcdA4 and Yp-TcaATcaB colored in a gradient from light to dark representing the pore domain with the TcB-binding domain, the α-helical shell, the β-sheet domains and the linker. The molecular model is shown for each TcAs and superimposed with the respective cryo-EM density. The colors correspond to those in Figure 1.

**Supplementary Video 2:** Pl-TcdA1 protomer with the molecular 3_1_ trefoil knot highlighted in rainbow colors corresponding to **Figure 3**. An alignment of the molecular knot structure of Pl-TcdA1 (red), Xn-XptA1 (blue), Mm-TcdA4 (yellow) and Yp-TcaATcaB (green) show the structural conservation of this region.

### Materials and Methods

#### Protein expression and purification

The expression and purification of the five TcAs were performed in a similar manner. The sequences of *tcdA1* (*P. luminescens*), *tcdA4* (*P. luminescens*), *xptA1* (*X. nematophila*) and *tcdA4* (*M. morganii*) were cloned in pET19d (Novagen). The sequences of *tcaa* and *tcab* (*Y. pseudotuberculosis*) were cloned together in pET28a (Novagen) with a GS-linker between the two fragments. All proteins were modified with an N-terminal hexahistidine tag.

Expression was performed in *E. coli* BL21 (DE3)RIPL. Cells were transformed with the specific plasmid and grown in medium (Xn-XptA1, Pl-TcdA1, Yp-TcaATcaB in LB medium, Mm-TcdA4 and Pl-TcdA4 in 2TY medium) until an OD_600_ of 0.6-0.8 at 37 °C. Protein expression was induced with 25 µM IPTG and performed at 18 °C for 15 hours (Xn-XptA1, Yp-TcaATcaB) or 20 °C for 20 hours (Pl-TcdA1, Pl-TcdA4 and Mm-TcdA4).

Cells were lysed in lysis-buffer with 200 µM Pefabloc using a microfluidizer. Lysis buffers were either composed of 20 mM Tris-HCl pH 8, 150 mM NaCl, 0.5 % Triton-X for Xn-XptA1 and Yp-TcaATcaB, or of 50 mM Tris-HCl pH 8, 150 mM NaCl, 0.05 % Tween-20 for Pl-TcdA1, Mm-TcdA4 and Pl-TcdA4. Soluble proteins were separated from cell debris by centrifugation (38,000 rpm, 30 min, 4°C) and loaded on a 5 ml HisTrap FF column (GE Healthcare). The N-terminally His-tagged protein was eluted with 20 mM Tris-HCl pH 8, 150 mM NaCl, 0.05 % Tween-20 and 500 mM imidazole. Protein-containing fractions were pooled and dialyzed against 20 mM Tris-HCl pH 8, 150 mM NaCl, 0.05 % Tween-20. Subsequently, the proteins were further purified by size exclusion chromatography using a Superose 6 10-300 column (GE Healthcare) for Xn-XptA1, Pl-TcdA4 and Yp-TcaATcaB or a Sephacryl S400 column (GE Healthcare) for Pl-TcdA1 and Mm-TcdA4. *P. luminescens* TcdB2-TccC3 was expressed and purified as described previously^10,14^. Site directed mutagenesis of the wild-type Pl-TcdA1 was performed according to standard procedures to generate the following mutants: E158A-R1873A, D965A-R1971A E1086A-R1166A, H1202A and Y1168F-Y1205F. The Pl-TcdA1 mutants were expressed and purified like Pl-TcdA1(WT).

#### Negative stain electron microscopy

4 µL of the protein solutions (0.05 mg/ml) were applied on freshly glow-discharged copper grids (Agar Scientific; G400C) with an additional layer of 8 nm carbon film. After 40 s incubation time, the grids were blotted with Whatman No 4 filter paper and stained with 0.75 % uranyl formate. Images were recorded using a Tecnai G Spirit electron microscope (FEI) or a JEOL-JEM 1400 electron microscope operated at 120 kV and equipped with a TVIPS TemCam F416 detector.

#### Sample preparation for cryo-EM

Quantifoil 2/1 grids with additional 2 nm carbon layer were used for Pl-TcdA1, Xn-XptA1 and Mm-TcdA4. C-Flat 2/1 grids with a self-made additional carbon layer were used for Pl-TcdA4. Quantifoil 2/1 grids without an additional layer were used for Yp-TcaATcaB. All samples were flash-frozen in liquid ethane using a Cryoplunger CP3 (Gatan) at 25 °C and ∼ 90 % humidity. The grids were freshly glow-discharged before sample application. For Pl-TcdA1, 3 µl of 0.08 mg/ml protein were incubated for 20 s and blotted for 1.8 s before plunging. For Pl-TcdA4, 4 µL of 0.1 mg/ml were incubated for 40 s and blotted for 2 s before plunging. For Xn-XptA1, 4 µL of 0.1 mg/ml protein were incubated for 40 s and blotted for 2.2 s before plunging. For Mm-TcdA4, 4 µL of 0.1 mg/ml protein were incubated for 40 s and blotted for 2.5 s before plunging. For Yp-TcaATcaB, 4µL of 3.5 mg/ml of protein were applied and directly blotted for 2 s.

#### Data acquisition

All data sets were collected on a TITAN KRIOS transmission electron microscope (ThermoFisher) equipped with a spherical-aberration corrector and an XFEG. The data set of Pl-TcdA4 was collected at the National Center for Electron Nanoscopy in Leiden (NeCEN). The other data sets were collected at the Max Planck Institute of Molecular Physiology, Dortmund. The images were recorded on a Falcon II direct electron detector for Pl-TcdA1, Pl-TcdA4, Xn-XptA1 and Mm-TcdA4 and on a Falcon III for Yp-TcaATcaB using the automated data collection software EPU (FEI). Four images were acquired per grid-hole in all cases. Detailed parameters of data acquisition (total dose and number of frames) for each TcA are listed in Supplementary Table 1.

#### Data processing

All images for each data set were inspected manually and micrographs with bad ice quality or high drift were discarded. Frame alignment was performed using motionCorrV.2.1 ^32^ for Pl-TcdA4, motionCor2 ^33^ for Xn-XptA1, Mm-TcdA4 and Yp-TcaATcaB, and unblur^34^ for Pl-TcdA1. In total, unweighted full-dose images, dose weighted images with full-dose as well as low-dose images (15 e^−^/Å^2^ for Pl-TcdA4 and 25 e^−^/Å^2^ for Xn-XptA1 and Mm-TcdA4) were generated. Data processing was performed with the software package SPHIRE^16^. CTF estimation was performed with CTER^35^ on the unweighted full-dose images. With the CTF assessment and the drift assessment tool in SPHIRE, micrographs with high defocus or drift were sorted out. The Pl-TcdA4 data set was picked manually with EMAN2 boxer^36^. For the Pl-TcdA1, Xn-XptA1 and Mm-TcdA4 data sets, we used the automated particle software Gautomatch (http://www.mrc-lmb.cam.ac.uk/kzhang/) for particle picking. Initially, ∼2,000 particles were picked manually with EMAN2 boxer and used for a first 2D classification with ISAC^37^ to generate class averages as templates for Gautomatch. For Yp-TcaATcaB, crYOLO was used for automated particle picking^38^. 15 images were picked manually and used as training data for crYOLO. Particles were extracted from the dose weighted full-dose images. Initial and final particle numbers are listed in Supplementary Table 1.

2D classification was performed using ISAC. For all TcAs, 3D refinement (sxmeridien) and 3D sorting (sxsort3d, both implemented in SPHIRE) were performed with imposed C5 symmetry. We used the map of Pl-TcdA1 (EMD-2297)^10^ and filtered it to 30 Å as an initial model. For the data sets of Pl-TcdA4, Xn-XptA1 and Mm-TcdA4, particles from the low dose images were extracted and the last iterations of the final refinement were performed in continuing mode with the low-dose particles. Sharpening was performed with a soft mask in SPHIRE, and the resolution was calculated between the two independently refined half maps at 0.143 FSC criterion (Supplementary Figure S2 and Supplementary Figure S3). The final densities were filtered to the estimated final average resolution. Local resolution of the obtained maps was determined using sxlocres of the SPHIRE software package and the final density maps were colored accordingly to the local resolution in Chimera ^39^. Detailed parameters for each data set are summarized in Supplementary Table 1.

#### Model building

The sequences of the five TcAs were aligned using Clustal Omega^40^. Homology models were generated for Xn-XptA1, Mm-TcdA4, Yp-TcaATcaB and Pl-TcdA4 based on the Pl-TcdA1 crystal structure (PDB-ID: 1VW1) using MODELLER^41^. The homology models were used as starting point for building the atomic models and were initially fitted in the EM density map using rigid body fitting in Chimera^39^. Some regions showed already a good fit, especially the channel domain as well as the α-helical shell. Other parts, especially in the RBDs, showed no good agreement with the density maps. These domains were fitted separately into the corresponding density with flexible fitting using iModFit^42^. The single fitted domains were then merged together into a single model. Nevertheless, some regions still had to be build or remodeled *de novo* in Coot^43^. For *de novo* model building, residues were only built when we could clearly trace the backbone within the density, otherwise we deleted these residues or complete domains (RBD C for Xn-XptA1 and Mm-TcdA4 and residues 1140-1239 for Yp-TcaATcaB). We used the real space refinement tool of PHENIX^44^ and Rosetta relaxation^45^ to refine the models. The geometries of the final refined models were evaluated with MolProbity^46^ and the data statistics are summarized in Supplementary Table 1.

#### Structure analysis and visualization

UCSF Chimera^39^ was used for structure analysis, visualization and figure preparation. Structure-based sequence alignments were generated using the T-Coffee expresso server^47^. For surface representations, we used protonated proteins generated with the H++ server^48^ according to pH 4, pH 7 and pH 11. For visualization of the protein surface electrostatics, the electrostatic Coulomb potential was calculated ranging from −14 to 14 kcal/mol in Chimera. The surface hydrophobicity of the TcA channels was colored according to the Chimera tool “define attribute” with residue specific scores as described previously^49^. The conservation values were based on the sequence alignment and RMSD values on the structure based sequence alignment^47^. The pore diameters of Pl-TcdA1, Xn-XptA1 and Mm-TcdA4 were calculated with ChExVis^50^ and MOLEOnline^51^. The protein knot was identified and determined using the online server knotprot^17^.

#### Black lipid membrane experiments

The single channel conductivity was measured by black lipid membrane experiments. The membranes were formed using 1 % solution of diphytanoyl phosphatidylcholine in n-decane (Avanti Polar Lipids, Alabaster, AL). The instrumentational setup consisted of a Teflon chamber with two compartments which are connected by a small hole with a surface area of 0.4 mm^2^. The lipid solution is painted across the hole, resulting in membrane formation. After the membrane turned black, toxin was added to the cis side (the black side) of the chamber. The membrane current was measured with a pair of Ag/AgCl electrodes with salt bridges. The electrodes were switched in series with a voltage source and homemade current amplifier on the basis of a Burr Brown operational amplifier as described previously^10^. All measurements were performed with a membrane potential of 20 mV. For the measurements with different pH values, buffer containing 1 M KCl and 10 mM citric acid pH 4, 10 mM MES pH 6 or 10 mM CAPS pH 11 were used. The average single channel conductance was calculated from at least 70 pore insertion events. For the measurements at pH 4, 6 and 11, 3.7 pmol, 19.4 pmol and 5.6 pmol of Pl-TcdA1, 0.28 pmol, 16.6 pmol, 0.4 pmol of Xn-XptA1, 3 pmol, 28.1 pmol and 4 pmol of Mm-TcdA4, and 29.3 pmol, 3.6 pmol and 16.4 pmol of Yp-TcaATcaB were used, respectively.

#### Holotoxin formation

For the interaction studies with *P. luminescens* TcdB2-TccC3 and the different TcAs, the complex formation was induced by mixing 1 µM TcdB2-TccC3 and 0.5 µM of the different TcAs (pentamer concentration) and incubated for 1 hour at 4 °C. To remove unbound TcdB2-TccC3, size exclusion chromatography was performed afterwards using a Superose 6 increase 5/150 column (GE Healthcare). The retention fraction corresponding to the holotoxin with a total protein concentration of about 20 nM was then visualized by negative stain EM.

#### Intoxication assay

HEK293T cells (Thermo Fisher) were intoxicated with pre-formed holotoxin composed of the different TcAs (Pl-TcdA1, Xn-XpyA1, Mm-TcdA4 and Yp-TcaATcaB) and Pl-TcdB2-TccC3. 2 × 10^4^ cells in 400 µl DMEM/F12 medium (Pan Biotech) were grown adherently overnight and subsequently 0.5, 5 or 10 nM of chimeric holotoxin was added. Incubation was allowed to continue for 16 h at 37 °C before imaging. Experiments were performed in triplicate. Cells were not tested for Mycoplasma contamination.

#### Biolayer interferometry (BLI)

Affinities of Pl-TcdA1, Xn-XptA1, Mm-TcdA4 and Yp-TcaATcaB to Pl-TcdB2-TccC3 were determined by BLI using an OctedRed 384 (forteBio, Pall Life Sciences) and Streptavidin (SA) Biosensors. Pl-TcdB2-TccC3 was biotinylated in 20 mM HEPES-NaOH pH 7.3, 200 mM NaCl, 0.01 % Tween20 (labeling buffer) with Sulfo-NHS-LC-Biotin (Thermo Scientific) in a 1:3 molar ratio for 2 h at room temperature, followed by 16 h at 4 °C. Unreacted Biotin label was washed out using AmiconUltra 100 kDa cutoff concentrators by diluting the sample two times with a 10-fold volume of measurement buffer (20 mM Tris-HCl pH 8.0, 200 mM NaCl, 0.01 % Tween20) and re-concentrating back to the original volume.

Biotinylated Pl-TcdB2-TccC3 was immobilized on SA biosensors at a concentration of 40 µg/ml, followed by quenching with 5 µg/ml biotin. BLI sensorgrams were measured in three steps: baseline (300 s), association (20 s for Pl-TcdA1 and Xn-XptA1, 200 s for Mm-TcdA4 and 60 s for Yp-TcaATcaB, respectively), and dissociation (300 s for Yp-TcaATcaB and 200 s for the other TcAs). The sensorgrams were corrected for background association of the respective TcA on unloaded SA biosensors. On- and off-rates of binding were determined simultaneously by a global curve fit according to a 1:1 binding model. All BLI steps were performed in measurement buffer with additional 0.3 mg/ml BSA.

#### Incubation of TcAs with nanodiscs at different pH values

The prepore to pore transition of Pl-TcdA1(WT) and the mutants was induced by pH-shift from pH 8 to pH 11. For Pl-TcdA1(WT) and the three alanine mutants (Pl-TcdA1-E158A-R1873A, Pl-TcdA1-D965A-R1971A and Pl-TcdA1-E1086A-R1166A), 50 nM toxin pentamer was incubated with 300 nM nanodiscs MSP1D1-⊗H5-His with POPC (Cube Biotech) and then dialyzed against 20 mM CAPS pH 11, 150 mM NaCl over a period of 72 h. For the mutants Pl-TcdA1-H1202A and Pl-TcdA1-Y1168F-Y1205F, 0.3 µM toxin and 2 µM nanodiscs (MSP2N2-His with POPC for Pl-TcdA1-H1202A and MSP1D1-⊗H5-His with POPC for Pl-TcdA1-Y1168F-Y1205F from Cube Biotech) were incubated and analyzed for 72 h against buffer at pH 11. Xn-XptA1 (0.4 µM) was incubated with nanodiscs (4 µM) MSP1E3D1-His (Cube Biotech) with brain polar lipids (Avanti Polar Lipids) in the presence of 3 mM CaCl_2_ and dialyzed for 72 h against buffer at pH 11. For Mm-TcdA4, 0.1 µM toxin and 1 µM nanodiscs (MSP2N2-His with POPC) as well as BPL liposomes (ratio 1:4) was analyzed at pH 11 and pH 5, respectively. The Yp-TcaATcaB toxin (0.2 µM) was incubated with 2 µM nanodiscs and dialyzed against either pH 4.7 (MSP2N2-His with POPC) or pH 10.5 (MSP1D1-⊗H5-His with POPC). Pore formation was checked with negative stain EM.

#### Nano differential scanning fluorimetry measurements

The melting temperatures of the different TcAs were determined by nano differential scanning fluorimetry (nanoDSF) in a Prometheus NT48 (Nanotemper). All measurements were performed in 20 mM Tris-HCl, pH 8, 150 mm NaCl, 0.05 % Tween-20 using 100 nM TcA (monomer concentration) and a temperature gradient of 1°C/min over a temperature range of 20-90 °C. For the pH stability test of Yp-TcaATcaB, a pH-range from 4 up to 11 was used with 20 mM citric acid for pH 4, 4.5 and 5, 20 mM Tris-HCl for pH 6, 7 and 8, 20 mM CHES for pH 9 and 10, 20 mM CAPS for pH 10.5 and 11 in addition to 150 mM NaCl and 0.05 % Tween-20 in all buffers. Measurements were performed in triplicates.

